# Optimization of dynamic soaring in a flap-gliding seabird and its impacts on large-scale distribution at sea

**DOI:** 10.1101/2022.03.03.482753

**Authors:** James A. Kempton, Joe Wynn, Sarah Bond, James Evry, Annette L. Fayet, Natasha Gillies, Tim Guilford, Marwa Kavelaars, Ignacio Juarez-Martinez, Oliver Padget, Christian Rutz, Akiko Shoji, Martyna Syposz, Graham K. Taylor

## Abstract

Dynamic soaring harvests energy from a spatiotemporal wind gradient, allowing albatrosses to glide over vast distances. However, its use is challenging to demonstrate empirically, and has yet to be confirmed in other seabirds. Here we investigate how flap-gliding Manx Shearwaters optimise their flight for dynamic soaring. We do so by deriving a new metric, the horizontal wind effectiveness, that quantifies how effectively flight harvests energy from a shear layer. We evaluate this metric empirically for fine-scale trajectories reconstructed from bird-borne video data using a simplified flight dynamics model. We find that the birds’ undulations are phased with their horizontal turning to optimise energy harvesting. We also assess the opportunity for energy harvesting in long-range, GPS-logged foraging trajectories, and find that Manx Shearwaters optimise their flight to increase the opportunity for dynamic soaring during favourable wind conditions. Our results show how small-scale dynamic soaring impacts large-scale Manx Shearwater distribution at sea.

**Teaser:** Flap-gliding shearwaters harvest wind energy by fine-scale trajectory optimization and this impacts their large-scale distribution at sea.

## Introduction

Wind speed and direction influence the movement ecology of birds across a range of spatiotemporal scales from local flight direction (*1*) and timing (*2–7*), through route selection (*6*) and drift compensation (*1, 8, 9*), to population distribution (*10*) and migration success (*5*). Such wind effects have been observed in a wide range of species, but the procellariiform seabirds (*10–14*), comprising the albatrosses, petrels and shearwaters, differ notably from non-pelagic species in their behavioral responses to wind. In contrast to the tailwind preference of non-pelagic birds (*2, 5–7*), Procellariiformes typically prefer crosswind or cross-tailwind flight (*11, 12, 14, 15*). They are also able to fly huge distances (*10, 16–18*) without having to offset their aerodynamic drag losses fully through flapping (*12, 19*). Both differences may be the product of a specialised flight mode called dynamic soaring (*20*), but whereas this is the only plausible explanation of how albatrosses are able to glide immense distances without flapping (*12*), dynamic soaring is more challenging to demonstrate in species such as shearwaters that routinely flap their wings during their characteristic rising-falling flight (*19*).

Soaring enables a bird to replace some or all of the aerodynamic kinetic energy that it loses to aerodynamic drag by harvesting energy from the atmosphere (*21*). Static soaring offsets these drag losses by harvesting gravitational potential energy from rising air currents, whereas dynamic soaring harvests aerodynamic kinetic energy from a spatiotemporal wind gradient (*22*). This can be most easily understood by noting that a bird flying into the wind experiences an increase in headwind speed as it moves from slower to faster moving air. Because the bird’s flight dynamics prevent its airspeed from equilibrating instantaneously (*23*), it follows that the bird will experience an immediate increase in its airspeed, and hence aerodynamic kinetic energy. Less obviously, perhaps, a similar increase in aerodynamic kinetic energy must also occur when a bird that is flying with the wind moves from faster to slower moving air. In general, dynamic soaring therefore entails flying *into* a strengthening wind and *with* a weakening wind (*22*). More specifically, because wind speed increases with height above the sea’s surface (*24*), pelagic dynamic soaring entails ascending *into* the wind and descending *with* the wind (*22*). We call these criteria the Rayleigh conditions (see Figure 1), after their late 19th century discoverer (*25*).

**Figure 1:**
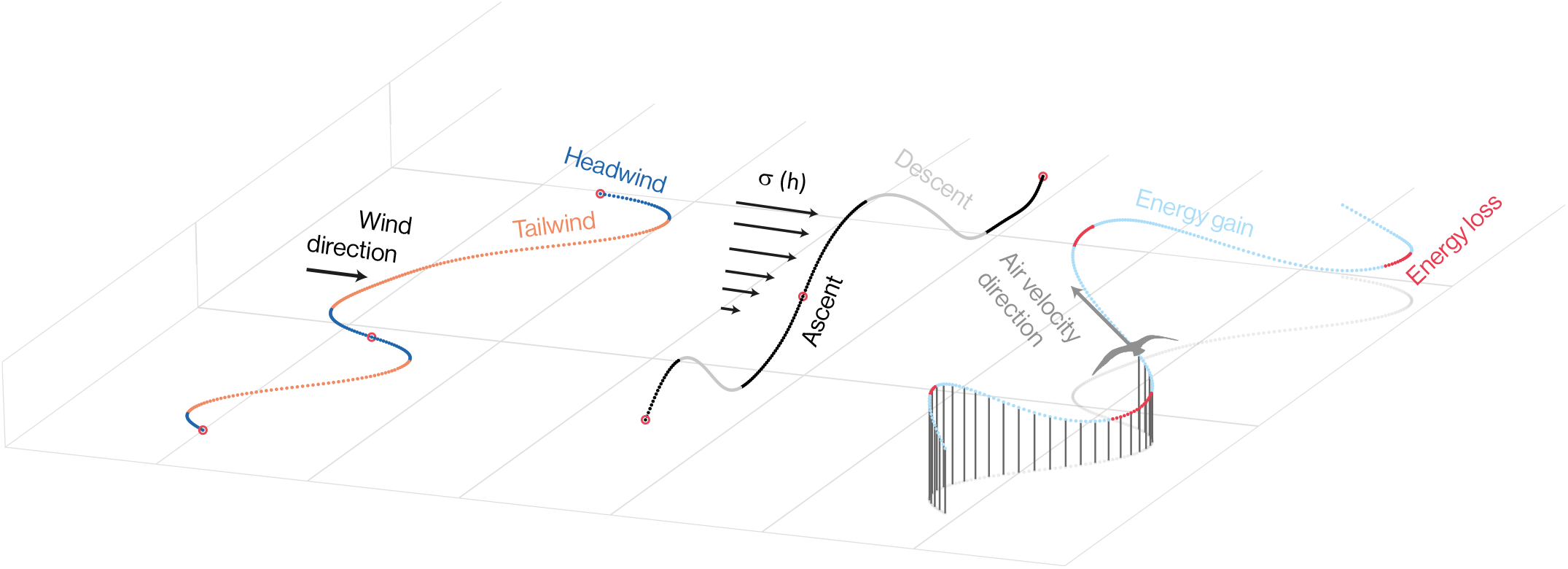
Cyclical basis of dynamic soaring. The energetic benefits of dynamic soaring are realised by phasing the horizontal and vertical components of flight to meet the Rayleigh conditions of flying into a strengthening wind and with a weakening wind. In a horizontal wind field, the sequence of headwinds (orange) and tailwinds (dark blue) that the bird experiences is determined only by the horizontal component of its flight trajectory (left). The vertical component (middle) determines whether the bird experiences an increasing or decreasing wind, according to whether it is ascending (black) or descending (light grey) through the shear layer (*σ*). The pattern of energy harvesting (right) depends on whether the phasing of the horizontal and vertical components of the flight trajectory meets the Rayleigh conditions, as these are what determine whether a bird gains (light blue) or loses (red) energy from the shear layer before taking account of drag losses. How much energy the bird harvests depends on the degree of alignment of the wind vector (black arrow) and the air velocity vector (grey arrow), as well as on the speed of flight and strength of wind shear. The trajectories shown here are reconstructed relative to the air from our empirical data for a part of one flight.

Although the physical principles of dynamic soaring are well understood, showing empirically that a bird is extracting aerodynamic kinetic energy from a spatiotemporal wind gradient—and hence implementing dynamic soaring—is extremely challenging. Among Procellariiformes, cyclical variation in mechanical energy has been used to assess dynamic soaring for the Wandering Albatross (*Diomedea exulans*) (*26*) and the Manx Shearwater (*Puffinus puffinus*) (*15*) using high-resolution GPS measurements. There is a general difficulty with this approach in that cyclical variation in mechanical energy does not necessarily imply cyclical dynamic soaring. For instance, a bird which is circling at constant airspeed in a uniform wind field displays cyclical variation in its inertial kinetic energy, but is not extracting energy from a spatiotemporal wind gradient. Similarly, intermittent bounding and undulating flight shows cyclical variation in both inertial kinetic energy and gravitational potential energy in the absence of energy harvesting from a spatiotemporal wind gradient (*27*). The mere occurrence of cyclical variation in mechanical energy is therefore insufficient to prove the use of dynamic soaring. In albatrosses, dynamic soaring has instead been inferred by analysing the distribution of mechanical energy gain with reference to a detailed aerodynamic model (*26*). Proving that flap-gliding species such as shearwaters use dynamic soaring is even more challenging, however, because the variation in mechanical energy that dynamic soaring entails may not even appear cyclical when changes in mechanical energy due to flapping are superimposed (*15*).

Here, we address these challenges by deriving a dimensionless metric describing dynamic soaring that we call the horizontal wind effectiveness (*ϵ*_*h*_). We estimate this empirically for flap-gliding Manx Shearwaters by using a simplified flight dynamics model to reconstruct *n* = 9 detailed flight trajectories from video data collected using bird-borne video loggers (*28*) attached to *N* = 6 individuals. The horizontal wind effectiveness *ϵ*_*h*_ quantifies how effectively flight in a given instantaneous direction harvests aerodynamic kinetic energy from a wind gradient, where positive values indicate that energy is gained from the shear layer and negative values indicate that energy is lost to it. Its long-term average 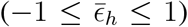 can therefore be used to assess whether and how a given flight trajectory is optimized for dynamic soaring in relation to the opportunity that the wind direction provides, which we quantify by defining another metric that we call the wind opportunity 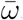. We find that shearwaters phase their vertical sequence of ascent and descent appropriately to gain energy from the shear layer independent of the wind speed *W*, but only optimize their horizontal turning cycle for dynamic soaring under favourable conditions when the crosswind component 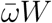 is high. We use GPS data from *n* = 368 outbound foraging flights and *n* = 368 return flights, from *N* = 201 individuals, to test whether this fine-scale behavior is recapitulated at larger spatiotemporal scales. We find that shearwaters tend to fly in a crosswind direction that is favourable for dynamic soaring on their outbound foraging trips, particularly when the wind speed *W* is high. Return legs are more constrained in their flight direction, and do not show any significant crosswind tendency. Besides conclusively demonstrating the use of dynamic soaring by a flap-gliding seabird, the metrics of wind effectiveness, wind opportunity, and crosswind component that we derive offer a framework for analysing dynamic soaring more generally.

## Theoretical Framework

### Energetics of dynamic soaring

For a bird flying in a steady horizontal wind field with vertical wind shear *σ ≥* 0, the mass-specific flow of useful mechanical energy due to dynamic soaring is shown in the Supplementary Text to be:

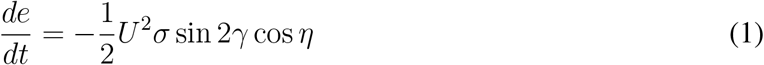

where *U* = ‖ **U** ‖ is the bird’s airspeed, *γ* is the aerodynamic flight path angle defined as the elevation of the bird’s air velocity **U** with respect to the horizontal, signed positive when climbing, and *η* is the heading-to-wind angle defined as the angle between the wind vector **W** and the horizontal component of **U**. The local shear gradient *σ* is usually unknown, but it is clear by inspection that the instantaneous rate of energy harvesting is always maximized by flying in an instantaneous direction that maximizes the dimensionless quantity:

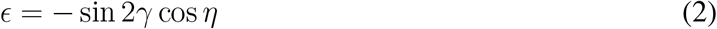

which we call the total wind effectiveness. Because wind velocity increases with height above a surface, *σ >* 0 by definition (see Supplementary Text). Energy is therefore harvested from the wind gradient when *ϵ >* 0 and lost when *ϵ <* 0, at a rate proportional to ‖*ϵ*‖.

The aerodynamic flight path angle *γ* is difficult to measure in an unknown wind field. However, because *−*90^°^ *≤ γ ≤* 90^°^ by definition, it follows that sin 2*γ* is positive when *γ >* 0, and negative when *γ <* 0. The instantaneous rate of energy harvesting is therefore maximized by flying in an instantaneous direction that maximizes the simpler quantity:

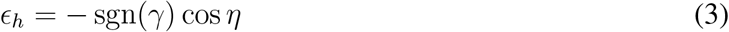

where sgn denotes the sign function. We call *ϵ*_*h*_ the horizontal wind effectiveness, because its magnitude depends only on the bird’s horizontal flight direction with respect to the wind *η*(*t*), whereas its sign depends on how the bird’s vertical motion *γ*(*t*) is phased with respect to its horizontal motion. Energy is harvested from the wind gradient when *ϵ*_*h*_ *>* 0 and lost when *ϵ*_*h*_ *<* 0, at a rate which is proportional to ‖ *ϵ*_*h*_ ‖. The Rayleigh conditions of ascending into a headwind (i.e. flying with *γ >* 0 and cos *η <* 0) or descending with a tailwind (i.e. flying with *γ <* 0 and cos *η >* 0) are met whenever *ϵ*_*h*_ *>* 0, but are implemented most effectively as *ϵ*_*h*_ *→* 1. Eq. 3 therefore emphasises that both the shape of the horizontal flight track and the phasing of its ascent-descent sequence are critical to determining how effectively energy is harvested through dynamic soaring.

### Flight dynamics model

Computing the horizontal wind effectiveness *ϵ*_*h*_ requires knowledge of the sign of the aerodynamic flight path angle *γ*, and measurement of the heading-to-wind angle *η* (see Eq. 3). Identifying the sign of *γ* is straightforward if the wind field is horizontal, because then sgn(*γ*) is positive when ascending and negative when descending. We will further assume that sgn(*γ*) = sgn(*θ*) where *θ* is the bird’s pitch angle (see Materials and Methods). Measuring *η* is much harder because whereas a bird’s flight trajectory is most readily obtained in an Earth-fixed coordinate system using a GPS logger, the heading-to-wind angle *η* is necessarily measured with respect to the air. Relating these two frames of reference requires knowledge of the local wind velocity, which is problematic in a shear layer. To avoid this difficulty, we instead use a simplified flight dynamics model to predict the turning associated with the bird’s pitching and banking motion that we measure by observing the horizon in bird-borne video from a body-fixed camera (Figure 2A,B; see also (*29*)). For this, we specify the bird’s orientation using a set of intrinsic Euler angles where *ψ* is the bird’s yaw angle, *θ* is the bird’s pitch angle, and *ϕ* is the bird’s bank angle, defined in that order (see Supplementary Text and Figure 2C–E).

**Figure 2:**
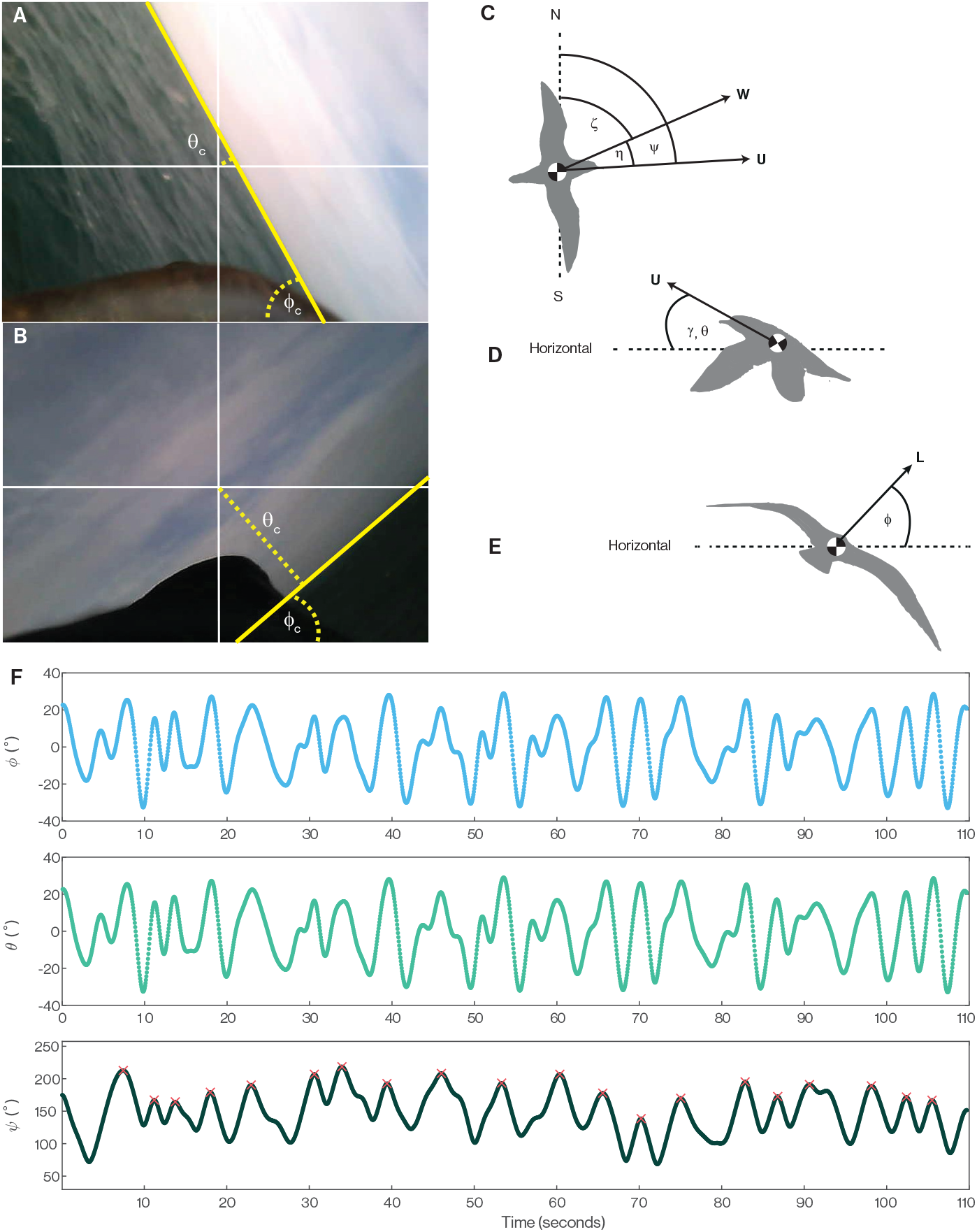
Reconstruction of cyclical turning behavior using bird-borne video. **(A**,**B)** Video from a camera mounted on the back of a Manx Shearwater demonstrates alternating left (A) and right (B) banked turns, coupled with cyclical pitching. The camera bank angle (*ϕ*_*c*_) is measured as the angle of the horizon line, which we detect automatically (yellow line); the camera pitch angle (*θ*_*c*_) is measured as the angular distance of the detected horizon line from the image centre (crosshairs); see Materials and Methods. **(C-E)** Angle definitions, shown in three orthogonal views. Overhead view (C): the bird’s heading-to-wind angle (*η*) is defined as the angle between the horizontal wind vector (**W**) and the bird’s air velocity vector (**U**) in horizontal projection; this angle *η* is assumed to equal the angular difference between the bird’s yaw angle (*ψ*) and the wind’s azimuth (*ζ*). Lateral view (D): the bird’s flight path angle (*γ*) is defined as the elevation of **U** with respect to the horizontal; this angle is assumed to equal the bird’s pitch angle (*θ*), which defines the inclination of the lift vector **L** in the bird’s symmetry plane. Caudal view (E): the bird’s bank angle (*ϕ*) defines the inclination of **L** in the bird’s transverse plane. **(F)** Sample time histories of the estimated bank (*ϕ*), pitch (*θ*) and yaw (*ψ*) angles for flight b117ii-753. Red crosses denote peaks in *ψ*, corresponding to points where the sign of *ϕ* changes; these points are used to define the beginning and end of each turning cycle.

We assume that the heading-to-wind angle *η* is equivalent to the angular difference between the bird’s yaw angle *ψ* and the wind’s bearing *ζ*, such that *η* = *ψ − ζ* (Figure 2C). This amounts to assuming that the bird’s air velocity vector **U** is aligned with its longitudinal body axis at all times. We estimate *ψ* by integrating the following approximate model of unsteady banked turning (see Supplementary Text) on a per-frame basis using trapezoidal integration:

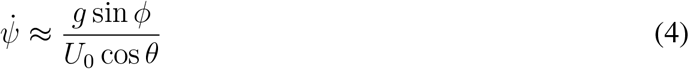

where *g* = 9.81 ms^-2^ is gravitational acceleration and *U*_0_ = 11.1 ms^-1^ is the bird’s equilibrium airspeed, as estimated in an earlier study of cruising Manx Shearwaters (*19*). Here, the angles *θ* and *ϕ* are the bird’s pitch and bank angles (Figure 2D,E), estimated from the apparent motion of the horizon (Figure 2A,B).

The unsteady banked turn model in Eq. 4 is an approximation, insofar as it assumes that the product of the bird’s wing area *S* and lift coefficient *C*_*L*_ remains approximately constant, and assumes that any variation in the airspeed *U* is small in comparison with the equilibrium airspeed *U*_0_. This is equivalent to assuming approximately constant airspeed *U ≈ U*_0_, and approximately constant lift *L* equal to body weight, *L ≈ mg* where *m* is mass. Eq. 4 also neglects the aerodynamic effects of wind shear on the bird’s turning, which is reasonable if the bird flies directly with or against the wind as it descends or ascends through the shear layer, as is most efficient for dynamic soaring (see Supplementary Text for detailed discussion). For the purposes of the subsequent statistical analysis, we define dynamic soaring cycles as beginning and ending at peaks in the estimated time history of the bird’s yaw angle *ψ* (Figure 2F).

### Optimization of dynamic soaring

Whilst it is possible in principle to find cyclical trajectories whose wind effectiveness is always non-negative, the optimization of a dynamic soaring trajectory is complicated in practice by the need to progress in a useful direction. The benefits of harvesting energy through dynamic soaring are therefore expected to trade off against the costs of deviating from the desired direction of progress, such that flights can typically be expected to involve alternating periods of negative as well as positive wind effectiveness. Because the rate of return from dynamic soaring is proportional to the wind shear strength *σ* (see Eq. 1), we should not necessarily expect birds to fly trajectories that maximize the energy they harvest on days with little wind. On such days, there is no prior reason to expect the mean wind effectiveness to be significantly different from zero, which is the null expectation if the bird phases the ascent-descent sequence described by *γ*(*t*) at random with respect to the pattern of horizontal turning described by *η*(*t*) (see Eq. 3). Conversely, under extremely windy conditions like those exploited by albatrosses flying in the Southern Ocean, it may be possible to harvest more energy than is lost to drag. Hence, whilst we should certainly expect the mean wind effectiveness to be positive under such favourable conditions, we should not necessarily expect it to be maximized outright, given that there is no way of storing any excess kinetic energy that may be harvested when flying close to the surface. In fact, energy harvesting is most likely to be maximized under marginally favourable conditions, when the returns available from dynamic soaring are sufficient to merit deviating from the desired direction of progress, but insufficient to enable flight to be sustained without flapping. These are precisely the conditions that we hypothesise will hold for Manx Shearwaters using flap-gliding flight on windy days.

## Results

Our bird-borne video data (Figure 2) show the weaving, undulatory, flap-gliding flight behavior that is characteristic of shearwaters (*19*). This is evidenced by the cyclical rolling and pitching of the visible horizon (Figure 2A,B), upon which are superimposed occasional faster oscillations due to flapping. The weaving component of this behavior is clearly visible in the *n* = 9 horizontal flight trajectories that we reconstructed at a fine scale by using Eq. 4 to model the turning associated with the birds’ observed rolling and pitching motion (Figure 3). Likewise, the coupling of the weaving and undulatory components of this behavior is apparent in the three-dimensional reconstructions (Figure 1). This turning behavior occurs on comparatively short timescales (median of mean turning cycle period per flight: 7.0 s), and produces an indirect flight path that must presumably offer some functional benefit to the bird. The reconstructed flight trajectories qualitatively resemble the cyclical trajectories attributed to dynamic soaring in albatrosses (*26*), and we analyse their suitability for harvesting energy from the shear layer quantitatively below. Nevertheless, as the shearwaters usually flapped their wings as they transitioned from ascending to descending flight, and as they were sometimes observed flapping at other points in the cycle, such energy as they harvested from the shear layer cannot have been sufficient to offset their drag losses completely. This fine-scale flight behavior is not directly observable in the coarse trajectories that we sampled at 5-minute intervals using GPS loggers, but it is reasonable to assume that the birds would have behaved similarly on these *n* = 368 flights, so we analyse the opportunity that these flights would have provided for dynamic soaring below.

**Figure 3:**
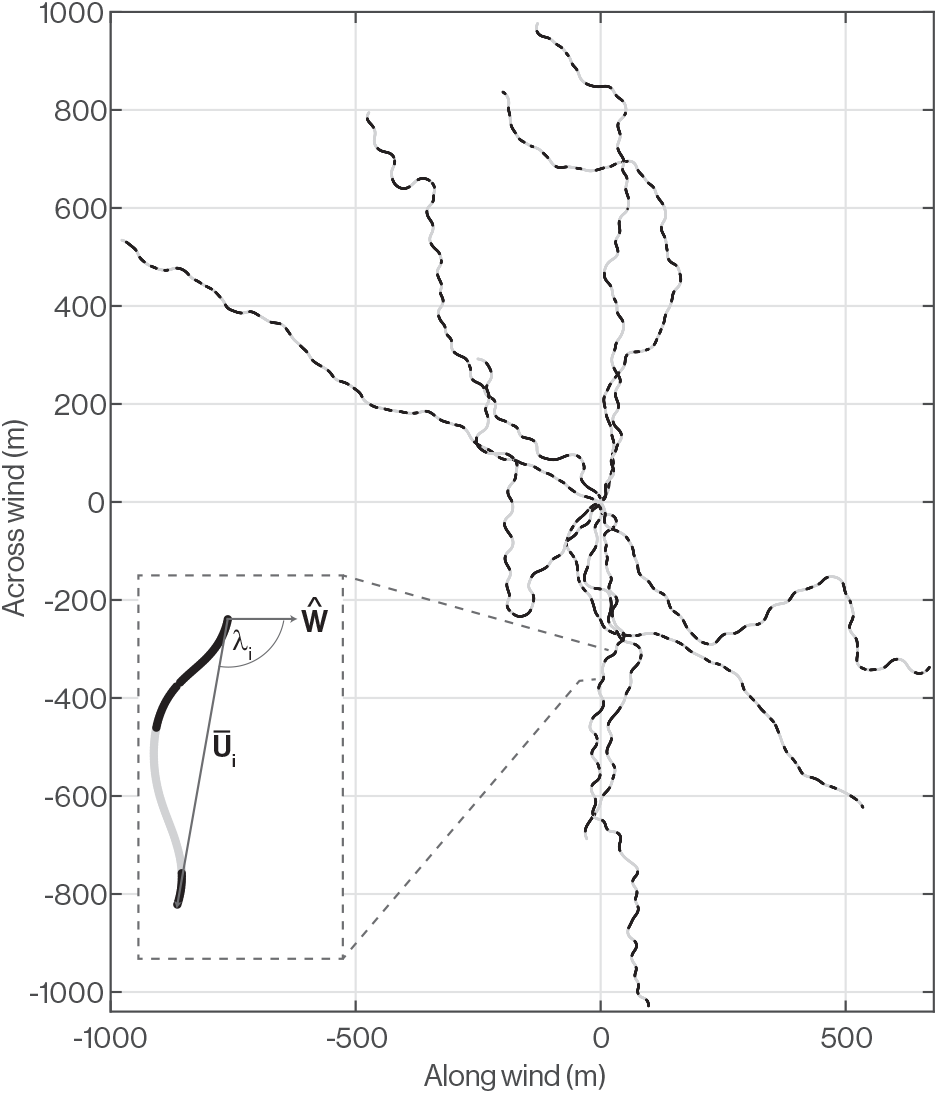
Weaving flight of Manx Shearwaters. All *n* = 6 individual Manx Shearwaters that we sampled at fine-scale consistently undertook weaving flight. We reconstructed their *n* = 9 trajectories shown here by using our estimates of the bird’s pitch, bank and assumed airspeed to estimate the bird’s air relative flight velocity vector **U**, which we integrated numerically using trapezoidal integration for the purposes of visualisation. Here, these air-relative trajectories are expressed in a coordinate system aligned with the experienced wind direction, such that the overall direction of each flight on the graph indicates the extent to which the bird heads with, against, or across the wind. The angle between the average air velocity vector Ū*_i_* of cycle and the wind velocity unit vector **Ŵ** we call *λ*_*i*_, and we use this for computing the mean wind opportunity metric 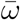 in section. Each trajectory has been projected into the horizontal plane and sections of ascent are shown in black, while sections of descent are shown in grey.

### The weaving flight of shearwaters implements dynamic soaring

The sample mean horizontal wind effectiveness 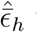 was positive on all *n* = 9 fine-scale flight trajectories that we collected from *N* = 6 individuals (sign test: *p* = 0.004, *n* = 9; median 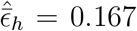 bootstrapped 95% CI: 0.042, 0.272). These data comprise a total of 17,457 video frames corresponding to approximately 873 s of flight data, with each recording lasting between 89 s and 114 s. This positive mean horizontal wind effectiveness indicates that the shearwaters flew in a manner expected to harvest energy from the shear layer, but how effectively were their trajectories optimized for dynamic soaring?

To answer this question, it will prove convenient to decompose the mean horizontal wind effectiveness 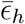 as:

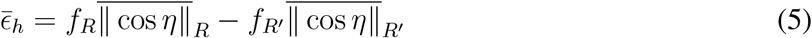

where *f*_*R*_ represents the frequency with which the Rayleigh conditions are met (i.e. the proportion of frames on which *ϵ*_*h*_ *>* 0) and *f*_*R*′_ = 1 *− f*_*R*_ represents the frequency with which they are not met (i.e. the proportion of frames on which *ϵ*_*h*_ *≤* 0). Implicitly, 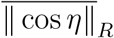 and 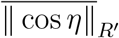 denote the mean magnitude of cos *η* over the corresponding portions of flight. This decomposition emphasises: (i) that the mean horizontal wind effectiveness 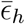 of a given pattern of turning behavior becomes maximal as *f*_*R*_ *→* 1, which entails phasing the ascent-descent sequence to meet the Rayleigh conditions; and (ii) that the maximal value of the mean horizontal wind effectiveness is further constrained by the time history of the bird’s horizontal flight direction with respect to the wind, because 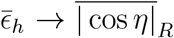 as *f*_*R*_ *→* 1. We will examine these two influences separately, first by assessing the optimality of the observed ascent-descent sequence conditional upon the observed horizontal trajectory, before then assessing the optimality of the observed horizontal trajectory in relation to the observed wind direction.

### Shearwaters phase their vertical and horizontal trajectories for dynamic soaring

We first compare the sample mean horizontal wind effectiveness 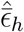 on each trajectory with the distribution of possible values of 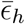 given counterfactual phasing of the same ascent-descent sequence. In other words, we compute the distribution of 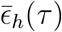 by holding the time history of the observed heading-to-wind angle *η*(*t*) fixed, while time-shifting the aerodynamic flight path angle *γ*(*t*) by an amount *τ* such that:

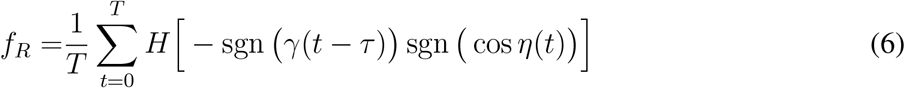

where *H* is the Heaviside function, which returns *H*(*x*) = 0 for *x ≤* 0 and *H*(*x*) = 1 for *x >* 0.

Likewise, we have:

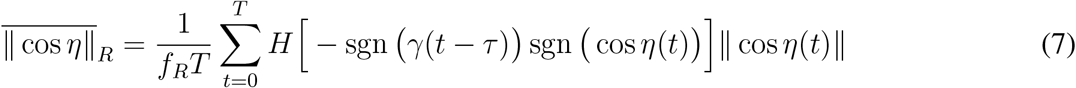

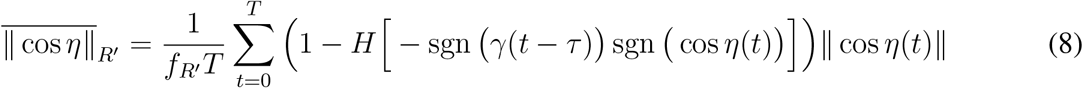

with *f*_*R*′_ = 1 *− f*_*R*_. Evaluating 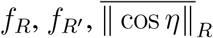 and 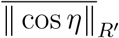 for different time shifts *τ* and substituting these into Eq. 5 generates the desired distribution of 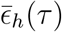, and recovers the observed value 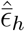 when *τ* = 0. Note that we wrap the time history of each flight when computing these quantities, such that the number of time points that we evaluate is the same for all values of *τ*. If the birds’ ascent-descent sequences were appropriately synchronised with their horizontal flight trajectories, then we would naturally expect 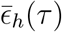 to peak at *τ ≈* 0. This was true of the ascent-descent sequences that we observed on all 9 flights (Figure 4), each of which was timed to within *±*0.4 s of the optimum, with a median phase shift of *−*7^°^ relative to the period of a complete turning cycle (1st, 3rd quartiles: *−*8^°^, 11^°^). Moreover, the sample mean horizontal wind effectiveness 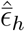 was usually close to the peak value of 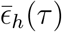, at a median value of 98.4% of the maximum (1st, 3rd quartiles: 94.6, 99.1%). These results demonstrate that the birds synchronised their ascent-descent sequences with their horizontal flight trajectories so as to harvest energy from the shear layer.

**Figure 4:**
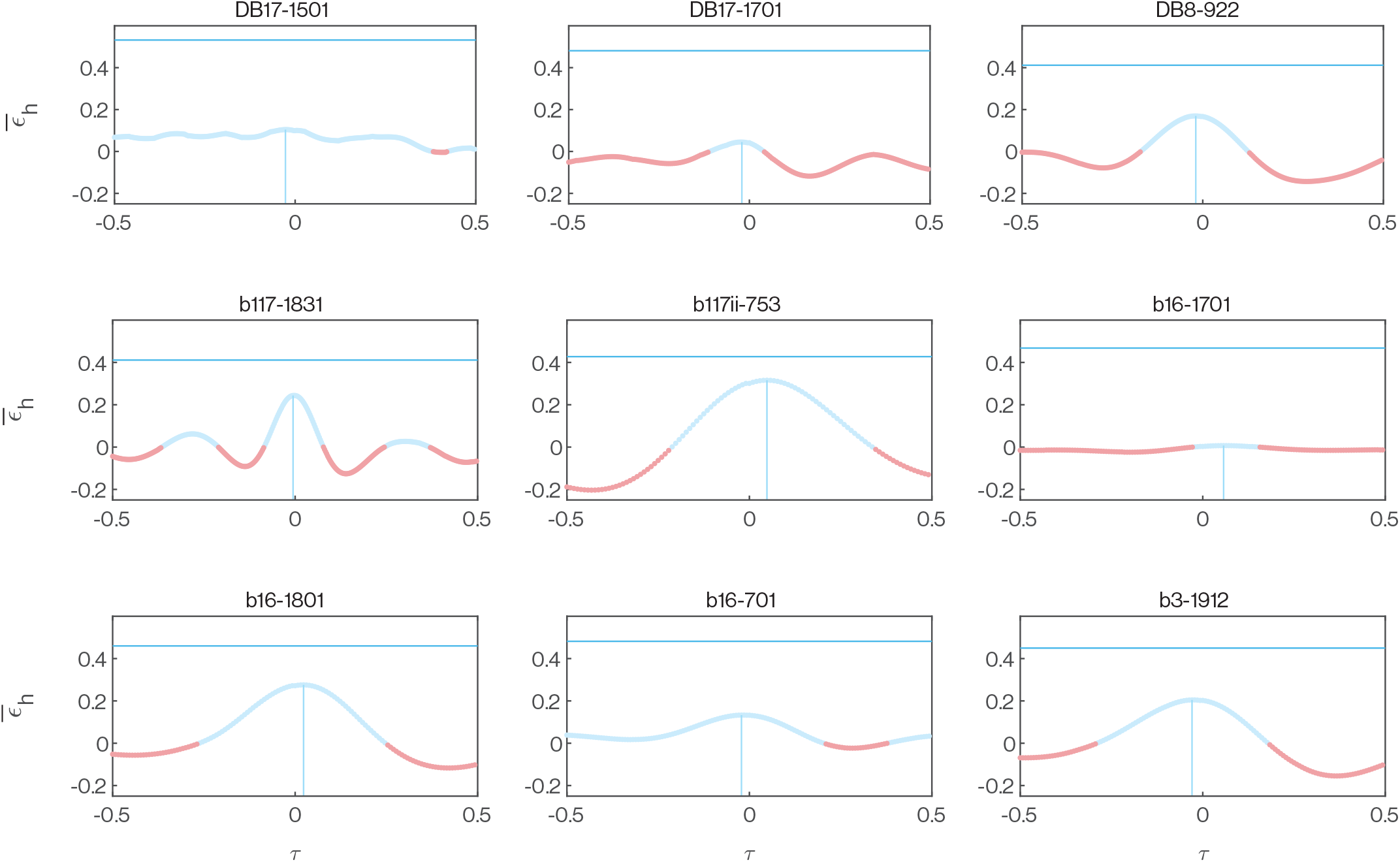
Optimization of the phasing of the vertical and horizontal components of each flight. Each panel plots the distribution of mean horizontal wind effectiveness 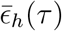 computed by holding the horizontal component of one of the *n* = 9 fine-scale flight trajectories fixed whilst varying the phasing of its vertical component by applying a time shift *τ*. Here −0.5 ≤*τ* ≤0.5, where *τ* is expressed relative to the mean period of one turning cycle. Positive values of 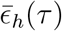 are shown in blue; negative values in red. The vertical blue line denotes the maximum value of 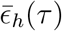 for each distribution and its associated value of *τ*, from which it is clear that the sample mean horizontal wind effectiveness 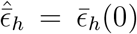 is close to the maximum 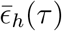 for each trajectory. The horizontal blue line plots the global optimum 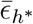 obtained by permuting the sequence of ascent and descent to maximize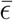, which is always substantially higher than the maximum 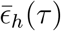 obtained by phase-shifting the ascent-descent sequence. See text for discussion.

This conclusion is based on holding the ascent-descent sequence fixed while varying its timing with respect to the observed horizontal flight trajectory. This is analogous to finding that a piano score harmonizes most closely when its bass and treble clef lines are correctly aligned, which provides evidence that they were jointly optimized in their composition, but says nothing about whether either is globally optimal. We therefore evaluate the maximum possible value that 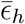 could have taken for each horizontal flight trajectory, by finding the optimal permutation of the observed ascent-descent sequence. That is, we find the global optimum 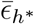, by permuting the observations of sgn(*γ*)(*t*) so as to: (i) maximize *f*_*R*_ at *f*_*R*_ = *f*_*R\**_ ; and (ii) maximize 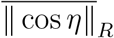 given *f*_*R*_ = *f*_*R\**_. This analysis showed that the sample mean horizontal wind effectiveness 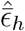 was considerably lower than the global optimum, at a median value of 40.5% of 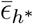 (1st, 3rd quartiles: 16.2, 59.2%)). We therefore find no evidence that the observed ascent-descent sequence was itself optimal, although it is worth noting that the strength of this conclusion depends on the fidelity of the modelled flight trajectory and assumed wind, and that there is no guarantee that any globally-optimal sequence of ascent and descent would have been physically achievable.

In principle, any sub-optimality of the observed ascent-descent sequence reflects the possibility of reordering it so as to: (i) meet the Rayleigh conditions more frequently; or (ii) meet the Rayleigh conditions at times when the horizontal flight direction was more effective for energy harvesting. To test between these alternatives, we compute a constrained global optimum 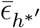, by permuting the observed sequence of sgn(*γ*(*t*)) so as to: (i) hold *f*_*R*_ fixed at 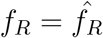, where 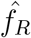 is the observed value of *f*_*R*_; and (ii) maximize 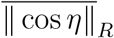 given that 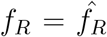. This analysis showed that the sample mean horizontal wind effectiveness 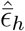 was considerably lower even than the constrained global optimum, at a median value of 56.6% of 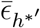 (1st, 3rd quartiles: 41.2, 74.9%), despite the Rayleigh conditions still being met with the same frequency 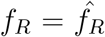. This result demonstrates that the sub-optimality of the observed ascent-descent sequences was primarily attributable to the Rayleigh conditions not being met at the most opportune points in the flight, and only secondarily to the frequency with which they were met. This is consistent with our earlier conclusion that the observed ascent-descent sequences were synchronised with the horizontal flight trajectories so as to harvest energy from the shear layer, but suggests that there might have been some room for improvement, subject to the earlier caveats on the fidelity of the modelled flight trajectory, assumed wind, and physical achievability.

### Shearwaters optimize their flight trajectories for dynamic soaring in a strong crosswind

The analysis so far has explored the optimization of the observed vertical sequence of ascent and descent conditional upon the observed horizontal flight trajectory, which is in turn constrained by the need to make useful progress over the sea’s surface. Theory predicts that a bird will maximize the energy it can harvest from the shear layer by heading in a net crosswind direction (*20, 30–32*). This is a natural consequence of the Rayleigh conditions, which state that energy is gained by ascending into the wind and descending with the wind, but lost if this phasing is reversed. Hence, because a bird must on average lose as much height as it gains to remain within the shear layer, any net progress parallel to the wind introduces an asymmetry that reduces the opportunity for energy harvesting. A bird may therefore increase the energy that it can harvest from the shear layer by biasing its flight direction to head across the wind on each cycle. We quantify this by defining the wind opportunity for the *i*th cycle as *ω*_*i*_ = ‖ sin *λ*_*i*_‖ where *λ*_*i*_ is the angle between the bird’s mean air velocity vector over that cycle **Ū**_*i*_ and the wind velocity unit vector **Ŵ** (Figure 3). In principle, a flight trajectory that heads across the wind on every cycle has a mean wind opportunity of 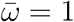, whereas a trajectory that heads into or against the wind on every cycle has a mean wind opportunity of 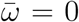. In practice, a value 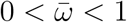 is expected if the bird changes its net heading across cycles.

To assess how the horizontal flight trajectories were optimized in relation to the observed wind direction, we compare the sample mean horizontal wind effectiveness 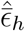 for each trajectory with the corresponding distribution of possible values of 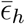 given counterfactual wind directions *ζ*. Specifically, we compute 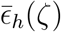 for each flight by holding the time history of the bird’s observed yaw angle *ψ*(*t*) fixed, whilst varying *ζ* in 1^°^ steps around the compass. This operation implicitly varies the mean wind opportunity 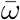 for each flight, which we use to assess the joint distribution of 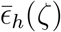 and 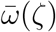. The resulting plots of 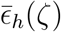 against 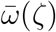 typically have a symmetric, scissor-like form comprising two looped arms (Figure 5). The symmetry of these plots reflects the fact that reversing the wind direction reverses the flow of energy, whereas the looped shape of each arm reflects the asymmetry of the birds’ turning cycles (Figure 3). In most cases, the maximum value of the wind opportunity 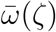 for any wind direction *ζ* was close to, but less than, one. This reflects the fact that the birds’ cyclical turning behavior was superimposed on a longer-term change in overall flight direction. Nevertheless, as we now show, the scissor-plots in Figure 5 provide clear evidence that the birds optimized their horizontal turning cycle for dynamic soaring on flights with higher crosswind components 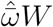. Conversely, on days when the crosswind component 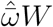 was lower, there is no clear evidence that the horizontal turning cycle was optimized for dynamic soaring—albeit that the vertical sequence of ascent and descent was still timed appropriately to gain rather than lose energy from the shear layer.

**Figure 5:**
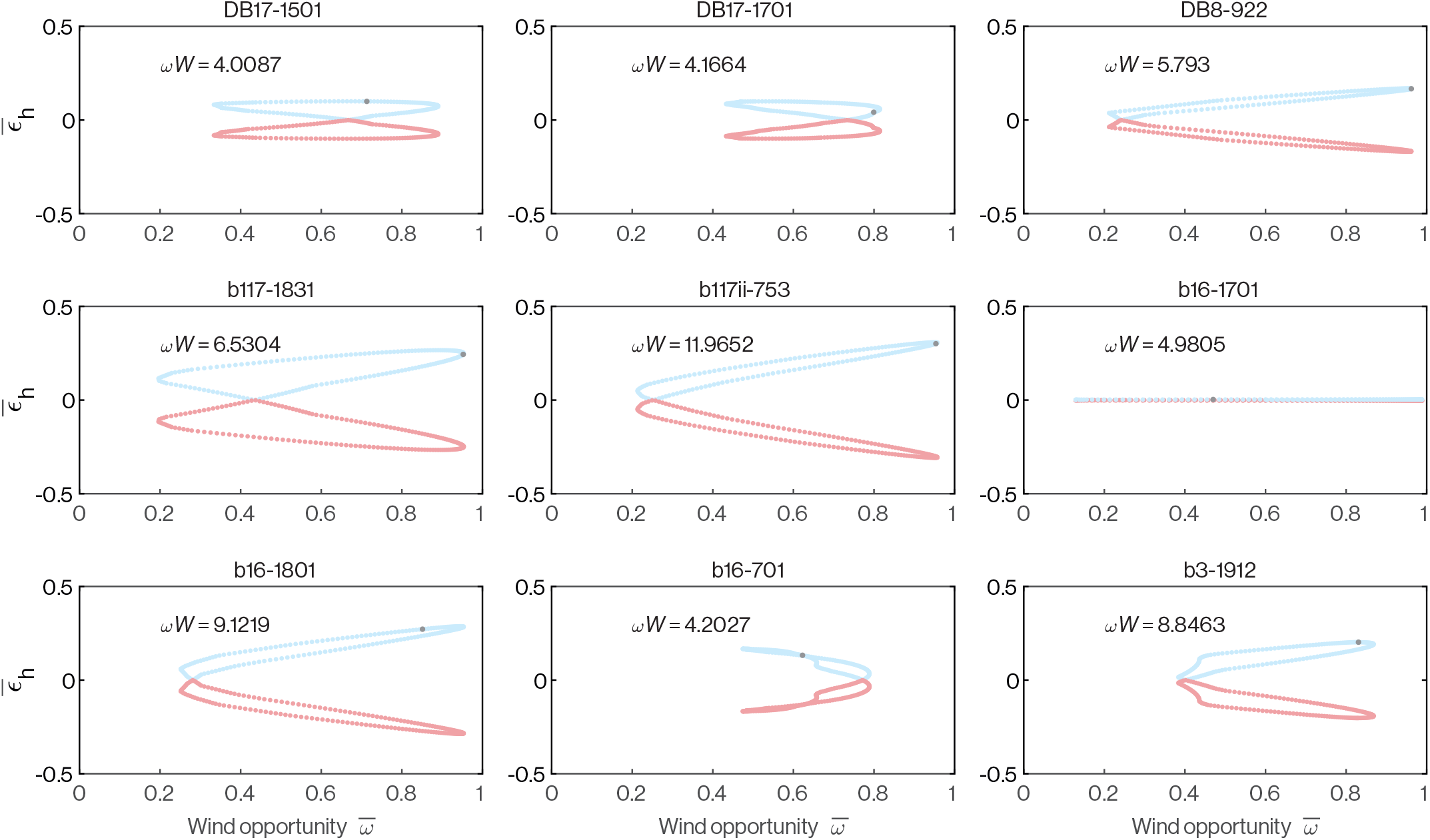
Optimization of horizontal flight trajectories with respect to the wind. Each panel plots the distribution of mean horizontal wind effectiveness 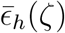 computed by holding the horizontal and vertical components of one of the *n* = 9 fine-scale flight trajectories fixed whilst varying the assumed wind direction *ζ*. Both the mean horizontal wind effectiveness 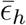 and the mean wind opportunity 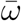 vary as the wind direction *ζ* varies, and plots of 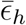 against 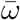 which have a scissor-like form in most cases. Positive values of 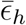 are shown in blue; negative values in red. The grey points mark the observed mean horizontal wind effectiveness 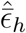 and observed mean wind opportunity 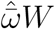. See text for discussion.

The proper phasing of the vertical sequence of ascent and descent to meet the Rayleigh conditions is evident from the fact that the observed point 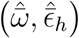 always lies on the upper arm of the scissorplot, for which 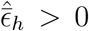 (Figure 5). Flights for which there is evidence that the detailed horizontal flight trajectory is itself optimized for dynamic soaring are those in which both of the following two conditions are satisfied. First, the maximum value of 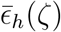 should coincide with a maximum of 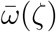. This implies that the turning cycle is tuned to harvest energy most effectively when the overall flight direction relative to the wind offers the most opportunity for energy harvesting. Second, the observed point 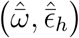 should be located at or close to the joint maximum of 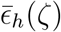 and 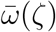. This implies that the observed flight trajectory has the greatest potential for harvesting energy when the wind direction is the same as was observed. These conditions hold for the 5 flights falling in the upper 50th percentile for the crosswind component 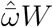, which are those offering the greatest potential returns from dynamic soaring. Conversely, these conditions do not hold on the 4 flights with crosswind components below the 50th percentile. In summary, the birds only turned in a manner that was tuned to capitalize on the opportunity for dynamic soaring on days when the potential return was high, albeit that they always phased their vertical sequence of ascent and descent to gain rather than lose energy from the shear layer.

### Shearwaters bias outbound flight in a crosswind direction favouring dynamic soaring

Given the strong association observed between mean horizontal wind effectiveness 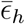 and mean crosswind component 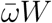 at fine spatiotemporal scales (Figure 6C), it is reasonable to expect that Manx Shearwaters might benefit from biasing the heading of their foraging flights across the wind, with possible implications for their large-scale distribution at sea. Depending on the locations of available foraging grounds, crosswind flight may not be possible in all winds on the outbound leg of a flight, and will be even more tightly constrained on the return leg by the unique direction to home. Indeed, pure crosswind flight is impossible to achieve on both the outbound and return legs of a flight, because directing the air velocity vector **U** perpendicular to the wind **W** on the outbound leg will generate a downwind drift requiring correction on the return leg. We might, therefore, expect to see a preference for crosswind flight directions on the outbound legs of foraging trips, which we test here against data from *n* = 368 outbound and *n* = 368 return tracks collected using GPS loggers attached to *N* = 201 individuals from 2016-2019 (Figure 7). Note that of the *n* = 368 tracks that we analyse for each direction of flight, 349 comprise the outbound and return legs of the same flights; the remaining 19 outbound and 19 return flights are not from matched samples.

**Figure 6:**
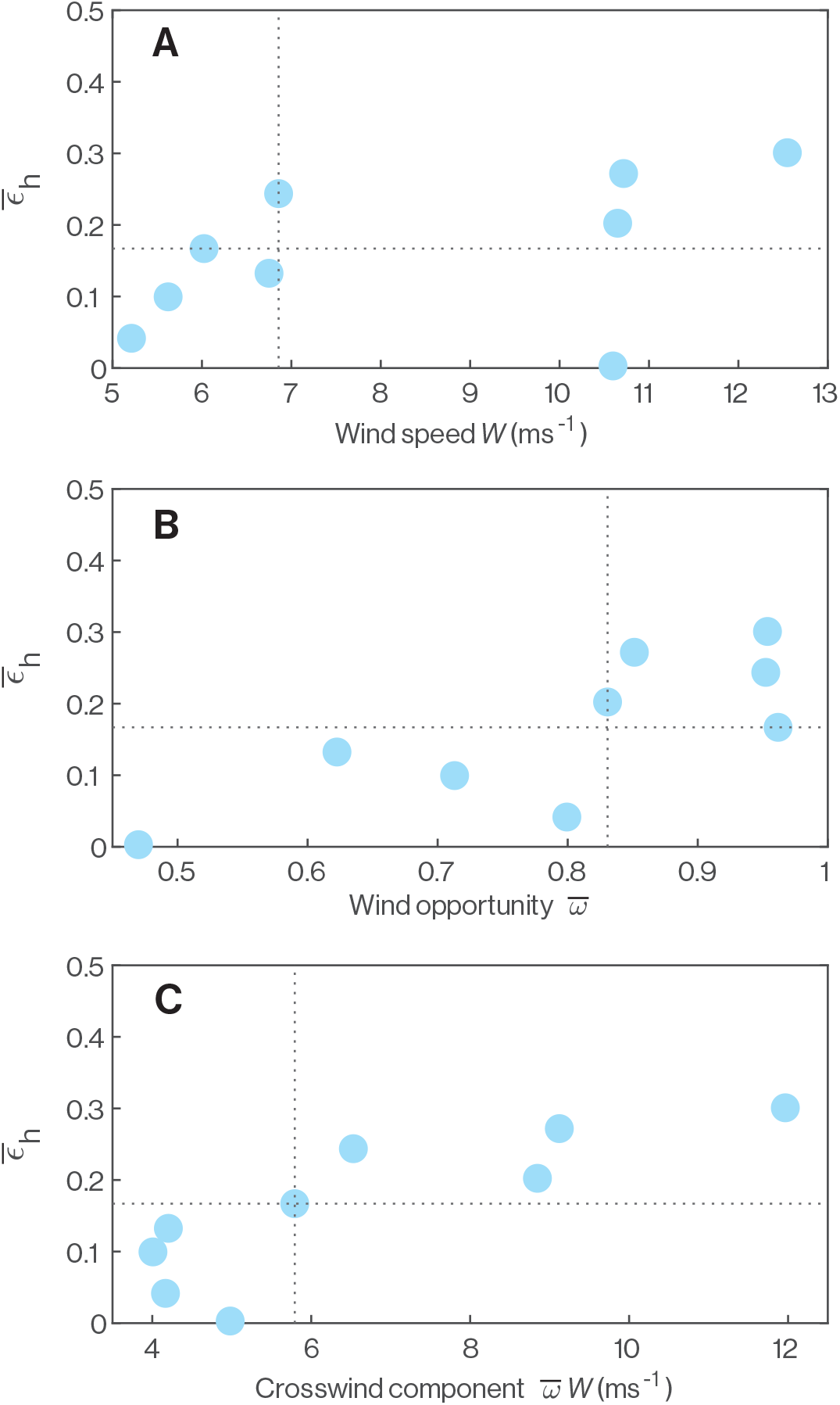
Influence of the wind on the effectiveness of dynamic soaring. Each datapoint represents one of the *n* = 9 fine-scale flight trajectories, and plots how the observed mean horizontal wind effectiveness 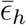 varies as a function of the wind and its interaction with the bird’s overall direction of flight. Wind speed *W* is treated as constant within a trajectory. The mean wind opportunity 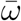 is the average of ‖ sin *λ*_*i*_ ‖ over a trajectory where *λ*_*i*_ is the angle between **W** and the bird’s average heading over the *i*th cycle. The crosswind component 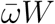 is the product of the wind speed *W* and mean wind opportunity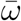. Vertical and horizontal dashed lines plot the median values of each variable. See text for discussion.

**Figure 7:**
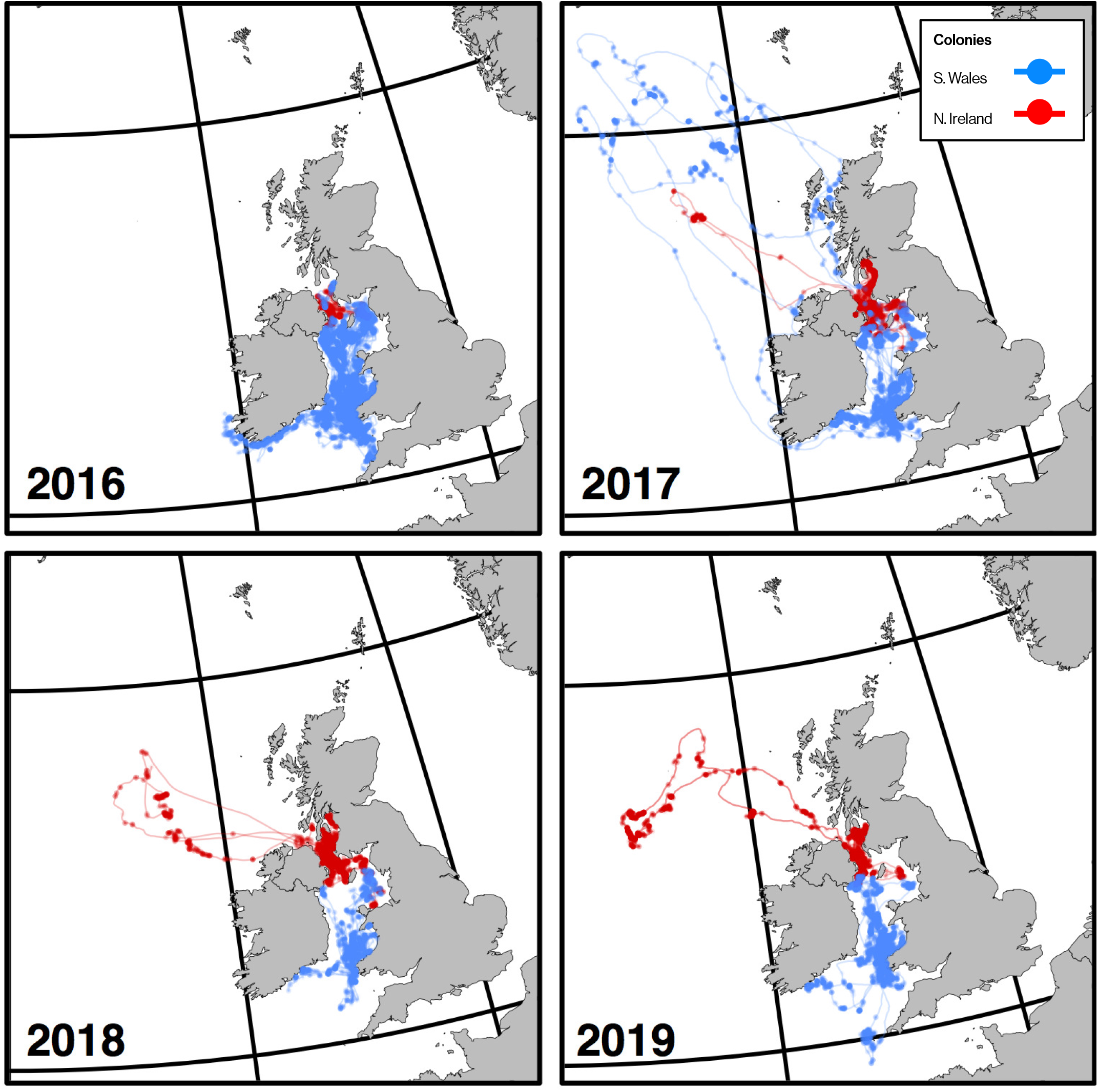
At-sea distribution of Manx Shearwaters. Coarse GPS trajectories show *n* = 368 flight tracks from colonies in Northern Ireland (Copeland; red) and southwest Wales (Skomer and Skokholm; blue) recorded from 2016-19. Feeding sites are highlighted with translucent points; sections of the track spent in directed flight are shown as connecting lines.

We calculated the overall heading-to-wind angle *λ* for each flight leg as the angle between the bird’s mean air velocity vector **Ū** over the flight leg and the wind velocity vector **W** (Figure 8A). As the benefits of heading in a crosswind direction are independent of whether the wind comes from the left or right, we quantified the variation in the overall heading-to-wind angle by computing the sample standard deviation of ‖ *λ* ‖ where *λ* is defined on the interval [*−*180^°^, 180^°^]. We then compared this to the population distribution of the same statistic obtained by randomising 100,000 times the pairings of the sample values of **Ū** and **W** that we used to calculate *λ* (Figure 8B). In this analysis we used all recorded outbound and return trajectories, meaning we included *n* = 368 outbound and *n* = 368 return trajectories (see Methods). We found that the overall heading-to-wind angle was no more conserved than expected by chance on the return legs of the flights (*n* = 368; *p* = 0.26), but was significantly conserved on the outbound legs (*n* = 368; *p <* 0.0001), albeit that the effect size was quite small (s.d. 50.1^°^ on return legs versus 48.2^°^ on outbound legs). The mean absolute heading-to-wind angle on the outbound legs was 81.3^°^ (*±*3.13^°^ 95% CI; Figure 8A), which is close to the crosswind tendency predicted to maximize the opportunity for dynamic soaring, but with a tailwind bias of 7^°^ appropriate to provide wind assistance on the outbound leg.

**Figure 8:**
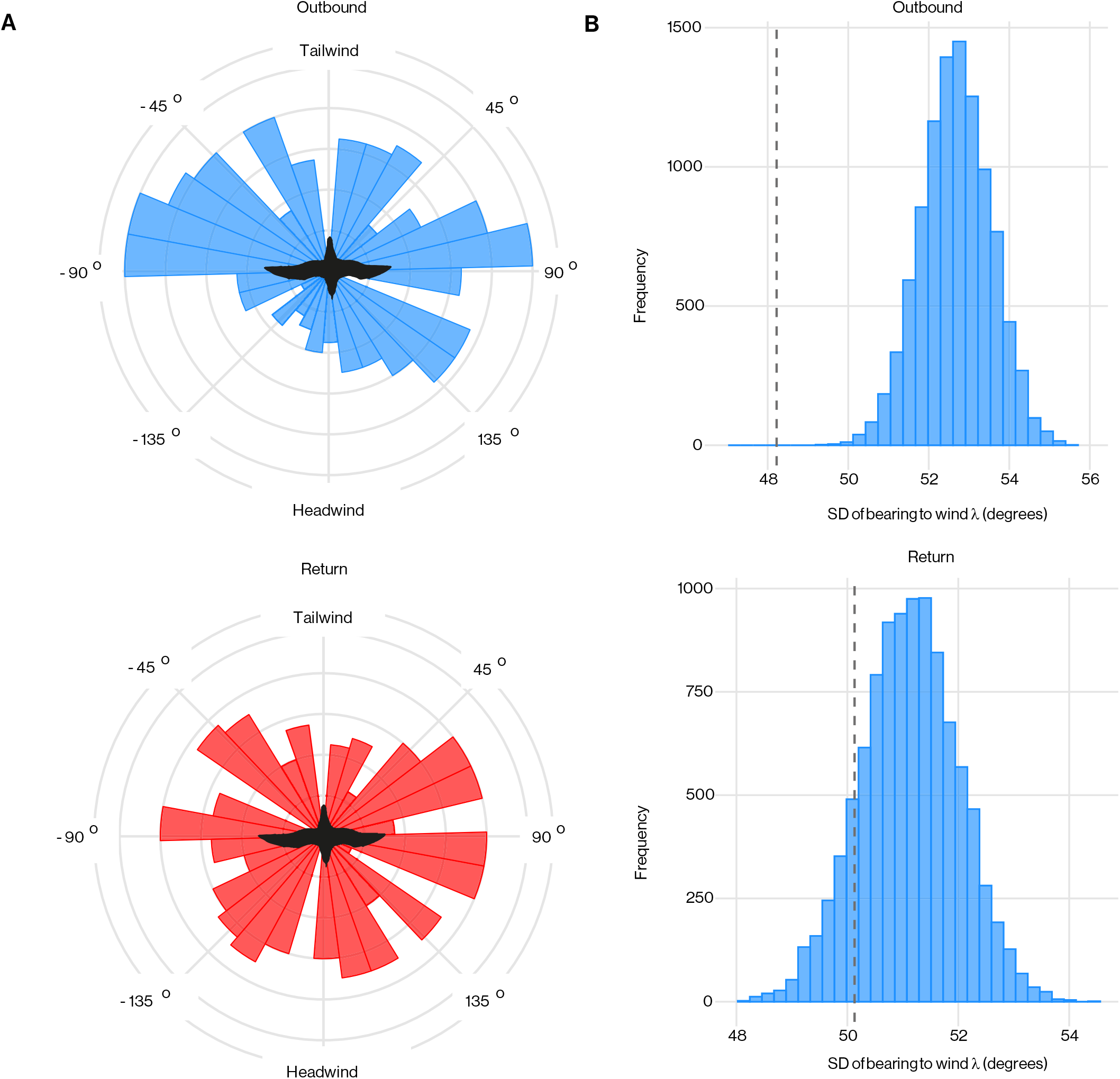
Association of wind direction and flight direction in Manx Shearwaters. **(A)** Circular histograms comparing the overall heading-to-wind angle (*λ*) experienced by shearwaters on outbound (top) and return (bottom) flight. **(B)** Distribution of standard deviation of absolute overall heading-to-wind angle ‖ *λ* ‖ computed by randomising the pairings of the sample flight directions and sample wind speeds on outbound (top) and return (bottom) flights; red dashed lines mark sample standard deviations.

To test how the observed crosswind tendency impacted the potential returns from dynamic soaring, we compared the mean wind opportunity 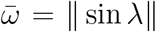 between the outbound and return legs of the flights. As expected, the mean wind opportunity 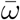 was significantly higher on the outbound journey for flights on which both legs were represented (*n* = 349; paired Mann-Whitney test; *p* = 0.0149), confirming that the birds’ tendency to fly across the wind on their outbound flights increased their potential returns from dynamic soaring (Figure 9). Furthermore, 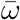 increased significantly with wind speed *W* on the outbound legs of the flights (Spearman’s *ρ* = 0.292; *p <* 0.0001; *n* = 368), but was not significantly associated with wind speed on their return legs (Spearman’s *ρ* = 0.0508; *p* = 0.331; *n* = 368). It follows that the birds’ tendency to fly across the wind on their outbound flights was stronger on windier days, when the potential returns from dynamic soaring were highest (Figure 9). This result could in principle have arisen as an artefact of the prevailing wind coming from a direction promoting crosswind flight. However, we found no evidence that the strongest winds came from a constrained direction when comparing the Rayleigh statistic for the wind directions associated with the top 10% of wind speeds with the population distribution of this statistic after randomising the pairings of wind speed and wind direction (*p* = 0.34). We conclude that the shearwaters’ tendency to forage in a more crosswind direction on windier days increases their potential returns from dynamic soaring when conditions are most conducive for harvesting energy from the shear layer.

**Figure 9:**
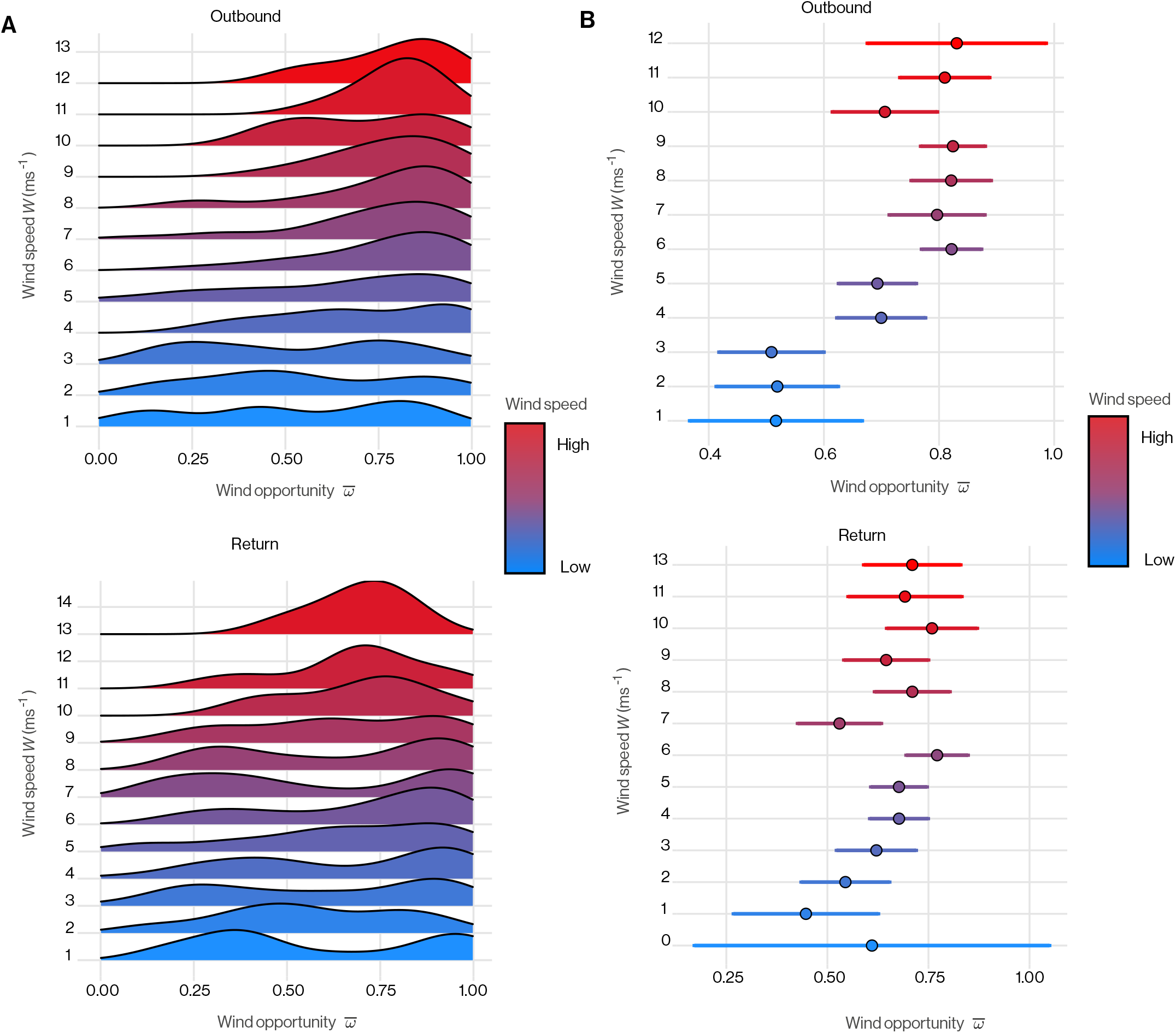
Association of wind speed and flight direction in Manx Shearwaters. **(A)** Density curves of mean wind opportunity 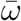 plotted for integer wind speed bins (m s^-1^). The higher the curve, the greater the density, and hence the more frequently that birds experience that wind opportunity when flying in winds of that speed. **(B)** Mean wind opportunity 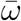 plotted with bootstrapped 95% confidence interval in integer wind speed bins (m s^-1^)

## Discussion

Here we use bird-borne video loggers to record the fine-scale flight trajectories of flap-gliding Manx Shearwaters, and use GPS loggers to record their coarse-scale movements and distribution at sea. Our data cover spatial scales ranging from less than a metre to hundreds of kilometres, and time scales ranging from less than a second to several days. We find that the shearwaters flew trajectories optimized for dynamic soaring, and that they harvested energy from the shear layer more effectively with increasing wind speed. Furthermore, we find that on outbound flights when their destination was less constrained, Manx Shearwaters increased the crosswind component of their flight trajectory to maximize the opportunity for energy harvesting, and did so to a greater extent when the wind speed was higher. We discuss below why such a strategy might be adaptive for these birds, and consider how small-scale flight decisions might influence large-scale feeding distribution at sea.

### Optimization of dynamic soaring in Manx Shearwaters

Pelagic birds harvesting kinetic energy from the shear layer incur an opposing cost because of the weaving and undulating flight trajectory needed to extract energy by dynamic soaring. This increases the path length relative to direct flight, and induces drag through the use of lift to turn (*23*). Dynamic soaring behavior therefore represents a trade-off between maximizing the potential for energy harvesting and minimizing the energy losses due to drag. Assessing both sides of this trade-off is challenging in flap-gliding species because of the difficulty of measuring flight efficiency in a bird using intermittent flapping. We have therefore focused on quantifying how effectively energy is harvested from the shear layer through the mean horizontal wind effectiveness and mean wind opportunity metrics that we have defined. We represent the associated trade-off diagrammatically in the form of a performance space (*33*) plotting mean horizontal wind effectiveness 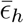 against mean crosswind component 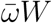 (Figure 10).

**Figure 10:**
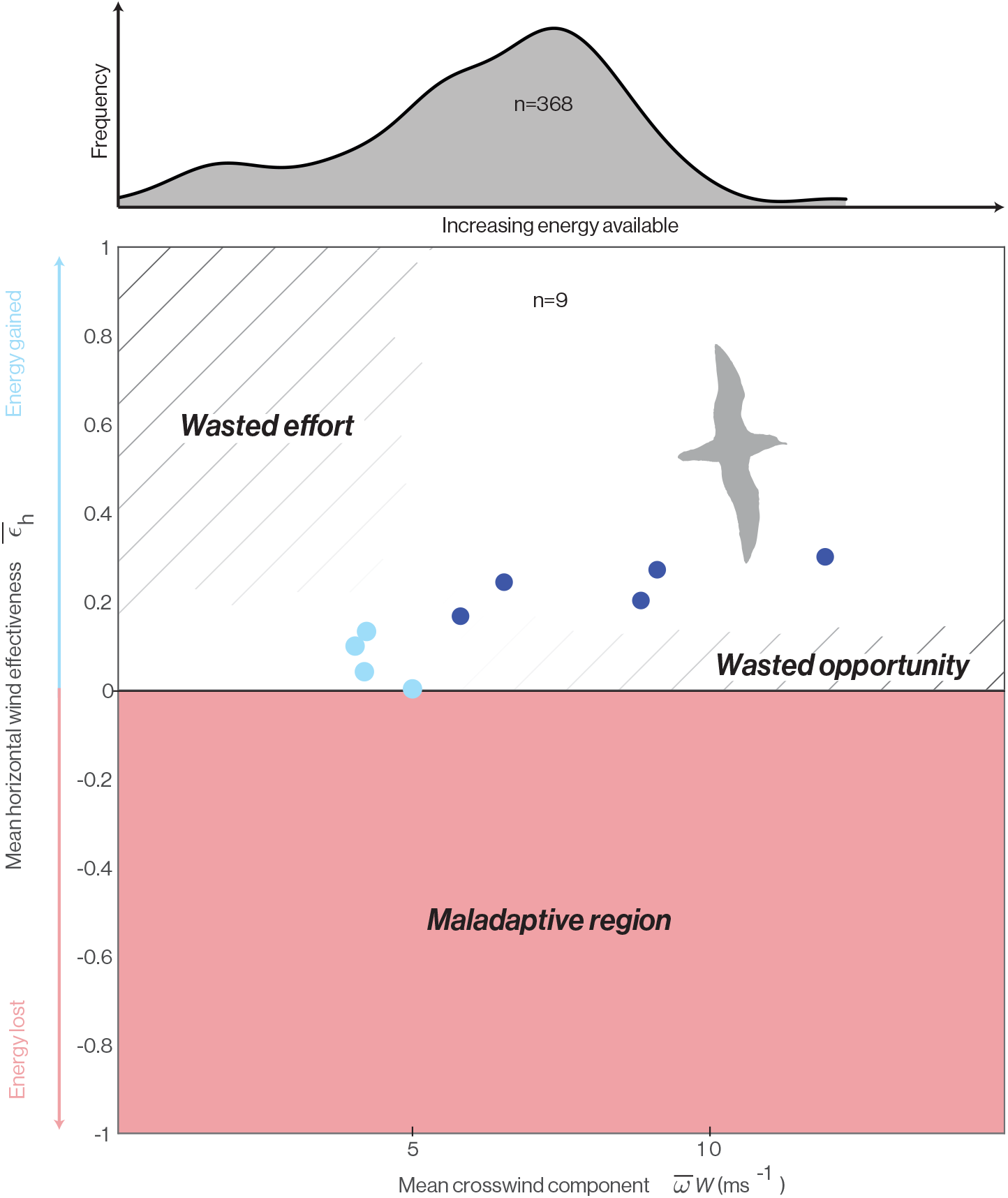
Performance space of dynamic soaring. Birds implementing dynamic soaring should fly such that they occupy the white region of the performance space. The red-shaded area is the region of performance space in which energy is lost to the wind gradient, which is maladaptive in the sense that it is always possible to find alternative ascent-descent sequences that avoid this. Within the upper half of the performance space, trajectories occupying the cross-hatched regions are sub-optimal in the sense that they represent either wasted effort or wasted opportunity. The wasted opportunity region lies to the right hand side of the performance space, as the energy accessible in the wind gradient scales increases with the mean crosswind component 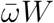. How this bears on our studied shearwaters is illustrated by the histogram at the top of the figure which shows the distribution of mean crosswind component 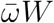 over the *n* = 368 recorded outbound flights. The blue-filled circles plotted in the performance space are the data from Figure (6C) where those shaded darker correspond to trajectories showing clear evidence of optimization of their horizontal trajectories in relation to the wind (see text for discussion). Note that the range of mean crosswind components over which there is clear evidence of detailed trajectory optimization for dynamic soaring (dark blue) matches the range experienced on the majority of flights (see histogram).

The lower half of this performance space (i.e.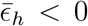) is unambiguously maladaptive, because flight in this region of the graph actively loses energy to the wind gradient (pink shaded region in Figure 10). Indeed, the shearwaters avoided this region of the graph on all *n* = 9 trajectories that we recorded at a fine scale (blue points in Figure 10). This region of the performance space represents a hazard for any bird ascending or descending through a shear layer, and we call this outcome maladaptive because it should always be possible to avoid. Specifically, for any horizontal trajectory, there will always exist corresponding ascent-descent sequences for which 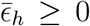. Unless there is some countervailing reason to ascend or descend, these should always be adopted in favour of ascentdescent sequences for which 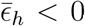. The reality of this hazard is demonstrated by the pink-shaded portions of Figures 4 and 5, which show how the same horizontal trajectory can be maladaptive given counterfactual phasing of its ascent-descent sequence or a counterfactual wind direction. Conversely, even the four flight trajectories experiencing the lowest mean crosswind component 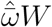, and hence the least opportunity for energy harvesting achieved 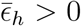 (Figure 10). The birds therefore avoided the small energy losses to the wind gradient that they would have experienced had they adopted poorly phased counterfactual ascent-descent sequences.

The upper half of the performance space (i.e.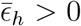) represents the region of the graph in which energy is actively harvested from the shear layer. However, because of the trade-off that exists between harvesting energy from the shear layer and losing energy to drag in the process, it is not equally beneficial to occupy all regions of this upper half of the performance space (Figure 10). Specifically, when the mean crosswind component of the trajectory 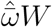 is low— whether because of low wind speed *W*, an unfavourable flight direction resulting in a low mean wind opportunity 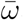, or both—there will be little energy accessible to harvest in the wind gradient. Increasing path tortuosity to maximize the effectiveness with which this little energy is harvested (i.e. maximizing 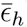) will not then yield appreciable increases in energy harvesting but will incur appreciable costs. The corresponding portion of the graph is therefore labelled ‘wasted effort’, referring to the crosshatched region in the top left of the performance space (Figure 10). The portion of the performance space labelled ‘wasted opportunity’ on the right hand side of the graph also represents a sub-optimal execution of the trade-off. In this region of the graph, the mean crosswind component 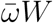 is high, so there is significant energy accessible to harvest in the wind gradient. Trajectories that do not effectively harvest this significant energy, as indicated by their low mean horizontal wind effectiveness 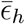, are therefore expected to be sub-optimal.

It is clear from Figure 10 that the shearwaters broadly conform to these expectations, avoiding the maladaptive region of the performance space altogether, and showing evidence of avoiding those regions of the performance space in which the most opportunity or effort is wasted. This implies that they synchronise their vertical motion appropriately with their horizontal motion so as to meet the Rayleigh conditions of ascending into the wind and descending with it. Nevertheless, using the fine-scale trajectory data that we derive from our bird-borne video footage, we only find evidence that the birds optimized their detailed horizontal trajectories in relation to the wind during flights on which the mean crosswind component 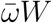 was high (Figure 5). These flights, marked as dark blue points in Figure 10, are those for which the evidence of dynamic soaring is strongest, so it is noteworthy that their mean crosswind components are in the same range as those recorded on the majority of the outbound flights that we sampled at a coarse scale using GPS (see histogram at top of Figure 10). In summary the GPS data show that for most of their outbound flights, the birds flew with a crosswind component that the video data show was associated with detailed trajectory optimisation for dynamic soaring.

While the value of the mean crosswind component 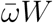 is obviously limited by the wind speed *W* served on the day, birds can maximize their mean crosswind component subject to this constraint by heading in a direction that maximizes the mean wind opportunity 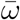. It follows that birds can move around the performance space in Figure 10 by making fine adjustments to their fine-scale flight trajectories to optimize their wind effectiveness 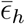, and by making coarse adjustments to their large-scale flight trajectories to optimize their mean crosswind component 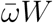 which implicitly alters 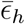. The performance space in Figure 10 thereby captures the effects of both fine and large-scale flight behavior.

Using the very large data set of *n* = 368 outbound flights and *n* = 368 return flights that we obtained using bird-borne GPS loggers, we find evidence that Manx Shearwaters tune their flights at this larger scale, by increasing the mean wind opportunity 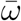 that they experience as wind speed *W* increases. In other words, shearwaters have a stronger tendency to fly in a crosswind direction on days when the wind is strongest (Figures 8), although we only observe this tendency during their outbound flights (Figure 9). This likely reflects a higher-level trade-off between adopting flight directions that are better for dynamic soaring and those that are better for other ecological reasons. In particular, flight direction is tightly constrained during homing flight because of the necessity to return to the breeding site to provision offspring. In contrast, constraints on flight direction are more relaxed during outbound flight for which the foraging destination is flexible within reason. It is important to note that the birds’ stronger tendency to fly across the wind on windier days could reflect headwind avoidance rather than crosswind selection. Nevertheless, as the effect of this large-scale behavior will be to promote dynamic soaring regardless, and as the birds’ fine-scale flight trajectories confirm their use of dynamic soaring, it seems reasonable to view the observed crosswind tendency as an adaptation for dynamic soaring on outbound flights.

How might the optimization of atmospheric energy harvesting impact the foraging distributions at-sea? Manx Shearwaters are known to utilise a dual foraging strategy when rearing (*34*), alternating between short chick-provisioning trips and longer self-provisioning trips to predictable oceanic resources (*35*). It is therefore possible that, within these constraints, decisions relating to choice of foraging site account for flight directions that increase energy extraction from the prevailing wind. Furthermore, given that the wind observed in the Irish Sea has a strong south-westerly bias, it is possible that the overall feeding distribution of shearwaters is contingent on these prevailing winds as a consequence of their use of dynamic soaring.

We have shown soft boundaries to the sub-optimal regions of the performance space to reflect uncertainty over their actual extent in relation to the performance that we observed in Manx Shearwaters (Figure 10). It is reasonable to assume that the performance spaces of other Procellariiformes will look broadly similar, but the upper half of the performance space might differ in detail for albatrosses, whose higher aspect ratio wings enable them to glide continuously over the windier open oceans. For example, for birds like the Wandering Albatross, we might expect the ‘wasted effort’ region to be narrower with respect to the mean crosswind component 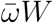. This is because the minimum mean crosswind component 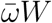 needed to fly energy-neutral dynamic soaring cycles is lower at the higher lift-to-drag ratios that higher aspect ratio wings enable (*36*). Moreover, at very high mean crosswind components, it is plausible that albatrosses might benefit from flying trajectories with reduced mean horizontal wind effectiveness 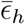 to avoid the very high aerodynamic loads associated with building up surplus kinetic energy.

### Implications for sensory physiology and behavior

Our finding that shearwaters optimise their fine-scale flight behavior for dynamic soaring might be thought to suggest that they require some sense of wind direction to determine when to transition between ascent and descent. In fact, for flight through a thick shear layer, it has been shown theoretically that optimal dynamic soaring trajectories approach a chain of alternating semi-circular turns (*23*). Under these conditions, a simple behavioral rule may suffice for the optimal control of dynamic soaring, because for a bird that spends as much time ascending as descending, transitioning between the two at the extremes of each cycle maximizes the energy gain (or loss) for any wind direction. All that is needed, therefore, is to select the binary phasing of ascent versus descent that makes the mean wind effectiveness positive, which only depends on knowing whether the wind is coming from the left or right of the average heading. The optimal transition points themselves are reached when the bird is heading in the same direction as its average heading, so could be sensed with the aid of any compass or other heading indicator, including a magnetic compass, the sun and its attendant cues, or distant landmarks and clouds (*37*). A detailed sense of wind direction would of course allow for more precise optimization of any trajectory, and might be possible using ventral optic flow cues under some conditions (*38*). In particular, if the bird’s longitudinal axis were aligned with its heading vector as assumed here, then any lateral optic flow must be a consequence of wind drift. Vision may therefore be at least as important for wind sensing in Procellariiformes as their tubular nostrils, which have been suggested to function as pitot tubes sensing airspeed during dynamic soaring (*39*).

Our camera-based method of reconstructing dynamic soaring trajectories also points to the fact that vision may be useful in controlling the turning flight behavior that we observe. When visible, the horizon offers a reliable means of identifying the vertical direction. This could be particularly useful in the control of pitching and banking during dynamic soaring, given that sensing gravitational acceleration is unreliable when aerodynamic accelerations are superimposed during unsteady manoeuvres.

In fact, our horizon sensing algorithm calls to mind the specializations of the visual field in Procellariiformes, which possess a horizontal streak where the retinal ganglion density is higher thereby improving resolving power around the horizon (*40*). Whilst this has traditionally been explained by the ‘terrain hypothesis’, which invokes the need for detection of objects against a horizon in open terrain (*41*) (*42*), it may also have a function in flight control (*43*). Specifically, by stabilising their head with respect to the fixed horizon, our shearwaters would have had an external reference for roll stabilisation from which, via proprioception, the angle of the body relative to the head could be inferred to obtain estimates of bank and pitch. This head stabilisation behavior is clearly visible in the videos. These sensing implications are also of interest for marine unmanned air vehicles (UAVs), given the potential to increase their flight time, range, and efficiency by dynamic soaring.

### Limitations

The horizontal wind effectiveness metric *ϵ*_*h*_ (Eq. 3) does not account for the quantitative effect of the bird’s flight path angle (*γ*) on the rate of energy harvesting. This is captured by the total wind effectiveness metric *ϵ* (Eq. 2), which differs from *ϵ*_*h*_ by a factor of sin 2*γ*. We have chosen to evaluate *ϵ*_*h*_ rather than *c* here on the pragmatic grounds that whereas it is usually possible to distinguish unambiguously whether a bird is ascending or descending, accurate quantitative estimates of the flight path angle *γ* relative to the air are difficult to obtain. Furthermore, the sign of the horizontal wind effectiveness *ϵ*_*h*_ is always the same as the total wind effectiveness *ϵ*, and evaluating the horizontal wind effectiveness metric *ϵ*_*h*_ partitions out the effects of the horizontal components of the flight trajectory, which are what combine to determine the overall flight direction that we analyse on larger spatiotemporal scales.

The mean horizontal wind effectiveness metric 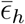 is calculated as the unweighted mean of the horizontal wind effectiveness *ϵ*_*h*_ across a flight trajectory. This is appropriate if a given flight direction is equally good at harvesting energy everywhere in the wind field, but this only strictly holds for a linear wind profile in which the vertical wind gradient is constant. In contrast, theoretical studies of dynamic soaring have variously assumed linear (*44*), exponential (*30*) (*31*) (*45*) (*46*), or logarithmic (*30*) (*31*) (*45*) models of wind shear. Under the nonlinear wind shear models, the wind gradient *σ* varies with altitude. For example, a logarithmic wind gradient is steepest just above the sea surface, and becomes shallower with height. In this case, the same flight direction and hence same value of *ϵ*_*h*_ would harvest more energy closer to the sea’s surface.

The suitability of the mean horizontal wind effectiveness metric 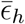 as a measure of how effectively a given flight trajectory harvests energy from the wind gradient therefore depends on the nature of the wind gradient, which is usually unknown. If necessary, the calculation of the mean horizontal wind effectiveness 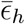 could be adjusted to account for the local wind gradient by taking a weighted mean of the horizontal wind effectiveness *ϵ*_*h*_ with weights that vary as a function of height. This operation requires precise knowledge of vertical position, however, which is difficult to achieve, even with GPS loggers. Hence, even in the presence of a nonlinear wind profile, there is utility calculating the mean horizontal wind effectiveness 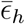 as we have done here, which only requires a binary assessment of whether the bird is ascending or descending.

Although the mean horizontal wind effectiveness 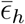 is itself a robust metric for assessing the effectiveness of a flight trajectory at harvesting energy from a vertical wind gradient, evaluating 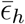 empirically requires accurate estimates of the flight path angle *γ* and yaw angle *ψ*. Clearly, the estimates that we have made of the bird’s pitch angle *θ* and bank angle *ϕ* are subject to measurement error, and although we have attempted to calibrate out any systematic bias in these estimates, our method for doing so is itself subject to model error (see Materials and Methods). Specifically, we derive the following exact expression for the yaw angle rate induced by an unsteady banked turn in a steady horizontal wind field of uniform direction with vertical wind shear *σ*:

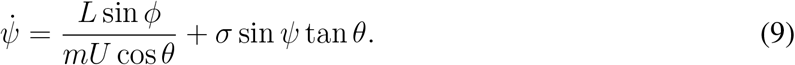

This expression is exact, subject to the simplifying assumption that the bird’s air velocity vector is always aligned with its longitudinal body axis (see Supplementary Text), which implies either that the bird is perfectly stable in pitch and yaw or that it coordinates its turning to achieve the same weathercock effect. For the purposes of estimating the bird’s yaw angle *ψ*, we have used the approximate model given in Eq. 4, which drops the term *σ* sin *ψ* tan *θ* describing the weathercock effect of wind shear. This is done on the basis that we know neither the vertical wind shear strength *σ*, nor the detailed stability properties of the bird that give rise to this term, but is expected to lead to model error (*33*) if the assumptions of Eq. 9 are satisfied.

Dropping this term is equivalent to assuming that the aerodynamic effect of wind shear on the birds’ turning is negligible, which is true when the vertical wind shear *σ* is weak, when the bird is flying with or against the wind such that sin *ψ* is small, or when the bird flies a shallow flight path such that tan *θ* is small. These conditions will not always prevail, resulting in a time-varying error in the estimation of 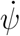. This will modify the shape of the weaving trajectories predicted under the model (Figure 3), but any changes in curvature will have a second-order effect on our estimates of mean horizontal wind effectiveness *ϵ*_*h*_ compared to any first-order integration error, which we remove by calibrating our estimates of *ψ* against the position of the sun’s disc in the video. In making this correction, we correct for camera bank offset, and we correct separately for camera pitch offset impacting our estimates of the flight path angle *γ*. Eq. 4 further assumes near-constant airspeed *U ≈ U*_0_, and near-constant lift equal to body weight *L ≈ mg*. Our constant airspeed assumption was necessary because we were unable to deploy airspeed sensors simultaneously with bird-borne video cameras given the constraints on carried load (*47*), but we do not expect this to have much effect on our results because airspeed tends to fall in a narrow range. We also find that any statistical tests reported to be significant at *U*_0_ = 11.1 m s^-1^ were also significant for alternative assumed airspeed values from 7.5 to 14.0 m s^-1^.

To evaluate the scope for behavioral sampling biases that could affect the generality of our findings, we used the STRANGE framework for animal behavior research (*48*). We used breeding birds for both GPS and video analysis so that loggers could be retrieved (*49*). Since both male and female shearwaters provide the same level of chick-rearing effort (*50*), it is unlikely our results are sex-biased. However, as breeding birds tends to be older than non-breeding birds, our samples will be age-biased. Given that flight behavior optimisation in relation to wind has been suggested to develop through learning (e.g. (*9*), it is therefore possible that our results might not generalize to younger, inexperienced shearwaters.

### A quantitative framework for identifying dynamic soaring

Although it is widely accepted that Procellariiformes use dynamic soaring (*14, 51*), and whilst the theory of dynamic soaring is well understood, rigorous empirical demonstrations of its utilisation are few and far between. Indeed, dynamic soaring has only previously been demonstrated conclusively in Wandering albatross (*26*), having been inferred somewhat indirectly in Manx Shearwaters (*15*). Demonstrating that a bird is implementing dynamic soaring demands showing that the bird is extracting energy actively from a wind gradient. One characteristic of such behavior is its cyclical variation in mechanical energy (see Introduction), but this is neither necessary nor sufficient to demonstrate dynamic soaring. It is not sufficient as mechanical energy may vary cyclically during other flight modes, such as the undulating intermittent flight of European Starlings (*Sturnus vulgaris*) (*27*). It is not necessary as mechanical energy variation due to drag losses, static soaring, and flapping (see Supplementary Text) may obscure the energetic signature of dynamic soaring altogether, and this is especially problematic in flap-gliding species such as Manx Shearwater (*15*). The metric of mean horizontal wind effectiveness 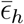 that we have derived here overcomes these limitations by assessing directly whether a given flight behavior is expected to harvest energy from the shear layer or to lose energy to it. Most importantly, this assessment is independent of other energy flows, such that it is not prone to masking in the way that an assessment of the changes in total mechanical energy would be.

Other observational studies have found no difference between large and small Procellariiformes in the frequency of flights that contain soaring (*51*), so our demonstration that Manx Shearwaters fly in a manner adapted to harvest energy from the shear layer suggests that dynamic soaring may be important across a broader range of species and environments than has been demonstrated to date. Demonstrating that the mean horizontal wind effectiveness 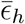 is consistently positive across flights, as we have shown here for Manx Shearwaters, therefore offers a reliable method for diagnosing dynamic soaring in other candidate flap-gliding Procellariiformes such as the Northern Fulmar (*Fulmarus glacialis*). When studied in combination with the other metrics of mean wind opportunity 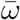 and mean crosswind component 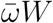 that we have derived here, the quantification of horizontal wind effectiveness also permits a fuller consideration of the optimization of dynamic soaring in these species. Furthermore, it could be applied to groups traditionally unassociated with dynamic soaring but known to glide in regions with strong spatiotemporal wind gradients such as Lesser Black-backed Gulls (*Larus fuscus*) flying in cityscapes (*52*), or Jackdaws (*Corvus monedula*) flying around ridgelines. Finally, our demonstration of dynamic soaring in coastal environments by a flap-gliding bird will be of interest to those working on marine unmanned aerial vehicles (UAVs). Our results illustrate the possibility of marine UAVs using dynamic soaring to decrease power consumption and thereby increase flight range, time and sustainability during facultatively powered flight in the socially important coastal zone.

## Materials and Methods

The theoretical framework that underpins the analysis described in the main text is derived in the Supplementary Text. The following sections describes the empirical methods used to collect and analyse the field data.

### Field observations

All work was done under ethical approval from the Animal Welfare and Ethical Review Board of the Department of Zoology, Oxford University, under endorsements from the British Trust for Ornithology Unconventional Methods Panel (BTO permit numbers C/5311 (Tim Guilford) and C/6128 (Oliver Padget)), and following approval by the Islands Conservation Advisory Committee.

We captured 146 bird-borne video recordings from Manx Shearwaters *Puffinus puffinus* during fieldwork undertaken 15–28 August 2014 on Skomer Island, Pembrokeshire (51.74^°^*N*, 5.30^°^*W*). However, as the behavior of interest for this study was cruising flight, we excluded videos containing take-off, landing, and sea-sitting behavior. Likewise, as the sun was needed to calibrate flight direction (see below), we excluded videos in which the sun’s disc was not visible, leaving a sample of *n* = 9 videos from *N* = 6 individuals across 6 different days.

Each video was captured using a miniature video logger (VEHO VCC-003 MUVI™ Micro DV Camcorders) (*28*) attached to the bird’s back using water soluble adhesive and TESA(4651) marine tape (*49*). A custom-designed, microprocessor-controlled timer was used to record two-minute sections of video once every hour (640 *×* 480 pixels; RGB24 AVI format; 20 frames per second). Video loggers were inserted into waterproof heat-shrink plastic tubing (*49*), in which a small incision was made to enable the head of the camera to be positioned upright pointing forwards. The camera head was inserted into a purpose-built Perspex turret, which was waterproofed and held in place by hotmelt glue. The video loggers weighed 17-18 g including battery, which is approximately 4% of body mass (*19, 47, 49*).

Shearwaters were also (not simultaneously) tracked at a coarse-scale using iGotU (Mobile Action, Taiwan) GPS devices (mass = 15g (*47*)) from colonies on Lighthouse Island (Copeland Archipelago; *n* = 164; 54.70°N, 5.52°W), Skomer island (*n* = 211; 51.74°N, 5.30°W) and Skokholm Island (2016 only; *n* = 10; 51.70°N, 5.30°W) from 2016 to 2019. These *n* = 368 tracks were taken from *N* = 201 birds in total. GPS loggers were set to record speed and position at 5-minute intervals, and were interpolated using a cubic spline function to ensure that the position estimates fell at precise 5-minute intervals (*53*). Tracks up to and including those gathered in 2016 were also analysed and published in (*54*).

### Video analysis

We estimated the camera pitch angle (*θ*_*c*_) and camera bank angle (*ϕ*_*c*_) directly from the video footage by writing an automatic horizon detection algorithm in MATLAB (The Mathworks Inc, Natick, MA). The algorithm squares the intensity of the blue channel to obtain a high-contrast grayscale image, to which it applies a Gaussian smoothing kernel with *σ* = 8. The algorithm then uses Roberts edge detection to find and mask the outline of the bird, and uses the Hough Transform to detect straight lines outside of this mask. The longest straight line is then taken as the putative horizon. We checked all putative horizon lines manually and edited these where necessary to correct any errors. The frame was skipped if no horizon line was visible.

We estimated the camera bank angle as *ϕ*_*c*_ = *−* arctan (Δ*y/*Δ*x*), where Δ*x* and Δ*y* denote the differences in the image coordinates of the horizon endpoints. Likewise, we estimated the camera pitch angle as *θ*_*c*_ = *s ×* 72*/*640, where *s* is the signed distance in pixels between the horizon line and the midpoint of the image, assuming an equiangular projection model over the measured 72^°^ field of view. We interpolated *ϕ*_*c*_ and *θ*_*c*_ for skipped frames using a piecewise cubic Hermite interpolating polynomial, and forwards-backwards filtered the signals using a fifth order low-pass Butterworth filter at a cut-off frequency of 0.5 Hz to remove high frequency noise due to measurement error, flapping perturbations, and other camera motion.

The measured pitch angle *θ*_*c*_ and measured bank angle *ϕ*_*c*_ of the camera differ from the true pitch angle *θ* and true bank angle *ϕ* of the bird because of camera offset and measurement error. We therefore estimate the bird’s pitch angle as 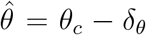 having calibrated the pitch offset *δ*_*θ*_ by enforcing the constraint that 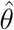 should average zero over the entire flight. This is justified on the basis that the vertical component of the bird’s motion *U* sin *θ* must average zero for it to remain close to the surface in a horizontal wind field, such that sin *θ* must also average zero if the airspeed *U* remains approximately constant as assumed.

We estimate the bird’s bank angle as 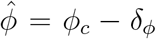, and calibrate out any bank offset *δ*_*ϕ*_ by integrating Eq. 4 numerically, having replaced *θ* and *ϕ* with their estimates 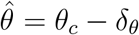 and 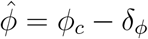. We estimated the bank offset *δ*_*ϕ*_ in piecewise fashion over a flight, by using the sun’s disc or a distant cloud as a reference for estimating the change in the camera’s azimuth Δ*ψ*_*c*_ between calibration frames at times *t*_1_ and *t*_2_, and finding the offset *δ*_*ϕ*_ that would satisfy the integral equation:

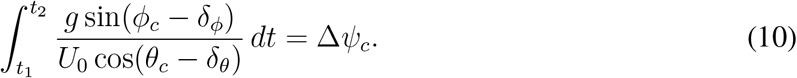

We measured the apparent azimuthal position of the sun in each calibration frame by measuring the signed distance *d* in pixels between lines dropped perpendicular to the horizon from the centre of the image and the sun’s disc or distant cloud. We then calculated the apparent change in azimuth Δ*ψ*_*c*_ between calibration frames at times *t*_1_ and *t*_2_ as Δ*ψ*_*c*_ = (*d*_2_ *− d*_1_) *×* 72*/*640 assuming the same equiangular projection model as before. This use of the sun as a compass serves to remove integration drift on each calibration interval, and also provides an external reference for relating the bird’s estimated yaw angle *ψ* to the wind direction *ζ* via the solar ephemeris. Specifically, we used the NOAA Solar Calculator to find the sun’s azimuth at the known date, time and approximate location of each video recording, from which we were able to calibrate the bird’s yaw angle relative to true north.

### GPS analysis

At-sea behavior was identified using a threshold model (*35*) combining information on the bird’s flight speed and the standard deviation of its heading recorded over a rolling window of 6 consecutive points. GPS fixes with a recorded ground speed of *>* 7*ms*^−1^ were classified as flight; those with a ground speed of *<* 7*ms*^−1^ and a rolling standard deviation in heading of *<* 18^°^ were classified as rafting; the remainder of points were classified as feeding. GPS fixes were only included in the analysis if they were classified as flight behavior.

The return sections of the GPS tracks were identified by moving backward along the track from the colony until the distance from the colony stopped increasing monotonically with respect to the length of the backward path (*55*). Conversely, the outbound section of the track was identified as the continuous section of the track over which the bird’s distance from the colony monotonically increased with total path length. To prevent pseudoreplication in the statistical analyses, the outbound and return sections of each track were summarised as single datapoints, with arithmetic mean values taken for wind speed and crosswind component, and with circular mean values taken for wind direction and heading in the air reference frame (see below). In total we included *n* = 348 flights for which both legs were tracked, and *n* = 38 where either the outbound or the return leg was recorded. As such, we include with each result presented in the main text a sample size that relates to the result in question.

### Wind measurements

Wind velocity was obtained from the NOAA/NCEP Global Forecast System (GFS). GFS data were sampled at a temporal resolution of 3 hours and a spatial resolution of 0.5^°^ *×* 0.5^°^. Each GPS fix was assigned the nearest wind measurement in time and space, which was used to determine the bird’s velocity with respect to the air as well as the ground. For the video-based analyses, the exact location of the shearwaters was unknown, but the wind velocity was found to be quite homogeneous within the known range of incubating shearwaters from Skomer (*56*) over the period for which the video footage was recorded. Given their known departure and arrival times, we estimate that the birds foraged within a 50 km radius of Skomer. We therefore averaged wind velocity estimates from the Met Office Wavewatch III wave model (one hourly temporal precision, 8km spatial precision, referenced to 10 m altitude) over this radius for the purposes of analysing the bird-borne video data.

## Acknowledgements

We thank members of the Oxford Navigation Group and Oxford Flight Group for insightful comments, and Glenn Wagner for help in deriving the model for an unsteady banked turn in a vertical wind gradient.

## Funding

University of Oxford Christopher Welch Scholarship (J.K.)

ASAB Undergraduate Project Scholarship (J.K.)

UKRI BBSRC scholarship grant number BB/M011224/1 (J.W., N.G.)

The Queen’s College, University of Oxford (A.F.)

Junior Research Fellowship at St. John’s College, University of Oxford (O.P.)

Merton College, University of Oxford (T.G.)

Mary Griffiths Award (T.G.)

BBSRC David Phillips Fellowship grant numbers BB/G023913/1 and BB/G023913/2 (C.R.)

Jesus College, University of Oxford (G.T.)

This project has received funding from the European Research Council (ERC) under the European Union’s Horizon 2020 research and innovation programme (grant agreement No 682501) (G.T.)

## Author contributions

Authors other than the joint-first authors are listed alphabetically. Individual contributions are as follows:

Conceptualization: J.K., J.W., T.G., G.T.

Formal analysis: J.K., J.W., G.T.

Investigation: J.K., S.B., J.E., A.F., N.G., T.G., M.K., I.M., O.P., C.R., A.S. and M.S.

Methodology: J.K., J.W., T.G., O.P., C.R., and G.T.

Supervision: T.G. G.T.

Visualization: J.K. and J.W.

Writing – original draft: J.K., J.W., G.T.

All authors reviewed and commented on the original draft.

## Data and materials availability

Raw GPS tracks are available at Movebank: doi:10.5441/001/1.k20j58qt and doi:xxx. Raw video data have been uploaded to figshare, together with the processed GPS and video data needed to replicate the analyses: doi:10.6084/m9.figshare.18019454.

## Supplementary Material

Supplementary Text

