## Supplementary Text for "Optimization of dynamic soaring in a flap-gliding seabird and its impacts on large-scale distribution at sea"

**Energetics of dynamic soaring** Here we derive an equation for the rate of change in useful mechanical energy during soaring. This equation demonstrates that one contribution is due to movement through a variable wind field, thereby quantifying the possibility for dynamic soaring. On the time-scale of our video recordings we can assume a flat, non-rotating Earth, such that the Earth-fixed axis system $\mathcal{F}_\mathcal{E} = (O, \hat{\mathbf{x}}_\mathcal{E}, \hat{\mathbf{y}}_\mathcal{E}, \hat{\mathbf{z}}_\mathcal{E})$ with arbitrary origin $O$ and right-handed orthonormal basis vectors $(\hat{\mathbf{x}}_\mathcal{E}, \hat{\mathbf{y}}_\mathcal{E}, \hat{\mathbf{z}}_\mathcal{E})$ with $\hat{\mathbf{z}}_\mathcal{E}$ pointing in the direction of gravity, is an inertial frame of reference. Denoting the centre of mass of the bird $G$, we call the bird's position vector in this coordinate system $\mathbf{r}_{G/O}$, such that its ground velocity $\mathbf{V}$ is the time-derivative of $\mathbf{r}_{G/O}$ in $\mathcal{F}_\mathcal{E}$, written $\mathbf{V} = {}^{\mathcal{E}}d\mathbf{r}_{G/O}/dt$. Likewise, if $\mathbf{r}_{W/O}$ is the position of an element of the surrounding fluid barring any influence of the bird, then the local wind velocity $\mathbf{W}$ is the time-derivative of $\mathbf{r}_{W/O}$ in $\mathcal{F}_\mathcal{E}$, written $\mathbf{W} = {}^{\mathcal{E}}d\mathbf{r}_{W/O}/dt$. Differentiating the identity $\mathbf{r}_{G/O} = \mathbf{r}_{W/O} + \mathbf{r}_{G/W}$ with respect to time in $\mathcal{F}_\mathcal{E}$, we therefore have:

$$\mathbf{V} = \mathbf{W} + \frac{{}^{\mathcal{E}}d\mathbf{r}_{G/W}}{dt} \tag{1}$$

which establishes $\mathbf{U} = {}^{\mathcal{E}}d\mathbf{r}_{G/W}/dt$ as the aerodynamic velocity of the bird, defined as the time-derivative of $\mathbf{r}_{G/W}$ in $\mathcal{F}_\mathcal{E}$. We may therefore rewrite Equation 1 as:

$$\mathbf{V} = \mathbf{W} + \mathbf{U} \tag{2}$$

The useful mechanical energy of a bird ($E$) is the sum of its aerodynamic kinetic energy and its gravitational potential energy, which we may write as:

$$E = \frac{1}{2}m(\mathbf{U} \cdot \mathbf{U}) - m\mathbf{g} \cdot \mathbf{r}_{G/O} \tag{3}$$

where $m$ is the bird's mass and $\mathbf{g}$ is the acceleration due to gravity. Hence, since $\mathcal{F}_{\mathcal{E}}$ is an inertial frame, the time-derivative of Equation 3:

$$\frac{{}^{\mathcal{E}}dE}{dt} = m\left(\mathbf{U} \cdot \frac{{}^{\mathcal{E}}d\mathbf{U}}{dt}\right) - m\mathbf{g} \cdot \mathbf{V} \tag{4}$$

gives the rate of change of useful mechanical energy in an inertial frame. Taking the derivative of Equation 2 in $\mathcal{F}_{\mathcal{E}}$ we have:

$$\frac{{}^{\mathcal{E}}d\mathbf{V}}{dt} = \frac{{}^{\mathcal{E}}d\mathbf{W}}{dt} + \frac{{}^{\mathcal{E}}d\mathbf{U}}{dt} \tag{5}$$

Then, by substitution of Equations 2 and 5 into Equation 4 , we have:

$$\frac{{}^{\mathcal{E}}dE}{dt} = m\mathbf{U} \cdot \left[\frac{{}^{\mathcal{E}}d\mathbf{V}}{dt} - \frac{{}^{\mathcal{E}}d\mathbf{W}}{dt}\right] - m\mathbf{g} \cdot [\mathbf{U} + \mathbf{W}] \tag{6}$$

such that by a rearrangement making use of the distributive property of the scalar product we have:

$$\frac{{}^{\mathcal{E}}dE}{dt} = \mathbf{U} \cdot \left[m\frac{{}^{\mathcal{E}}d\mathbf{V}}{dt} - m\mathbf{g}\right] - m\mathbf{g} \cdot \mathbf{W} - m\mathbf{U} \cdot \frac{{}^{\mathcal{E}}d\mathbf{W}}{dt}. \tag{7}$$

By Newton's second law, the first term of this equation is the total force acting on the bird less its weight, which is equivalent to the aerodynamic force $\mathbf{R}$. The vector equation for the total rate of change in useful mechanical energy can therefore be written as:

$$\frac{{}^{\mathcal{E}}dE}{dt} = \mathbf{U} \cdot \mathbf{R} - m\mathbf{g} \cdot \mathbf{W} - m\mathbf{U} \cdot \frac{{}^{\mathcal{E}}d\mathbf{W}}{dt} \tag{8}$$

The aerodynamic force $\mathbf{R}$ can be decomposed as the sum of aerodynamic lift $\mathbf{L}$, acting in a direction perpendicular to $\mathbf{U}$, and the net of aerodynamic drag $\mathbf{D}$ acting opposite to $\mathbf{U}$, and thrust $\mathbf{T}$ acting in the direction of $\mathbf{U}$. We therefore have $\mathbf{U} \cdot \mathbf{R} = (T - D)U$ where $T = \|\mathbf{T}\|$, $D = \|\mathbf{D}\|$, and $U = \|\mathbf{U}\|$ are the magnitudes of their respective vector quantities. Since $\mathbf{g}$ points down, only the vertical component of the wind vector ($W_z$) lies in its direction, such that $\mathbf{g} \cdot \mathbf{W} = gW_z$. Here $W_z$ is the component of wind in the direction of gravity and $g = \|\mathbf{g}\|$. We thus arrive at the simpler expression:

$$\frac{{}^{\mathcal{E}}dE}{dt} = \underbrace{TU - DU}_{\text{power surplus/deficit}} + \underbrace{mgW_z}_{\text{static soaring}} \underbrace{-m\mathbf{U} \cdot \frac{{}^{\mathcal{E}}d\mathbf{W}}{dt}}_{\text{dynamic soaring}} \tag{9}$$

which quantifies all of the energy flows relevant to soaring flight.

The flow of specific energy ($e$) due to dynamic soaring is:

$$\frac{{}^{\mathcal{E}}de}{dt} = -\mathbf{U} \cdot \frac{{}^{\mathcal{E}}d\mathbf{W}}{dt} \tag{10}$$

which is just the dynamic soaring term of Equation 9 normalised by the bird's mass $m$. Estimating ${}^{\mathcal{E}}de/dt$ empirically requires precise knowledge of the wind experienced by the bird, which is exceedingly difficult to measure in the natural environment. However, if the wind is assumed to have the same direction everywhere, given by the constant unit vector $\hat{\mathbf{W}} = \mathbf{W}/W$ where $W = ||\mathbf{W}||$ is the wind speed, then ${}^{\mathcal{E}}d\mathbf{W}/dt = \hat{\mathbf{W}}{}^{\mathcal{E}}dW/dt$. We will further assume that the wind is steady and horizontal, which implies that only the horizontal component of the bird's air velocity contributes to the scalar product in Equation 10. These assumptions are those generally employed in modelling dynamic soaring, and we may use them to rewrite Equation 10 as:

$$\frac{{}^{\mathcal{E}}de}{dt} = -U \cos\gamma \cos\eta \frac{{}^{\mathcal{E}}dW}{dt} \tag{11}$$

where $U = \|\mathbf{U}\|$ is the bird's airspeed, $\gamma$ is the aerodynamic flight path angle defined as the elevation of the bird's air velocity $\mathbf{U}$ with respect to the horizontal, and $\eta$ is the heading-to-wind angle defined as the angle between $\hat{\mathbf{W}}$ and the horizontal component of $\mathbf{U}$. If we further assume that $W$ is only a function of vertical height $h$ above the surface, where $\sigma(h) = {}^{\mathcal{E}}dW/dh$ is the vertical wind shear gradient, then we may make use of the bird's vertical climb rate ${}^{\mathcal{E}}dh/dt = U \sin\gamma$ to write the identity ${}^{\mathcal{E}}dW/dt = \sigma(h) U \sin\gamma$. Equation 11 then becomes:

$$\frac{{}^{\mathcal{E}}de}{dt} = -U\sigma(h)U \sin\gamma \cos\gamma \cos\eta \tag{12}$$

which we may rewrite using the trigonometric identity $\sin 2\gamma = 2 \sin\gamma \cos\gamma$ as:

$$\frac{{}^{\mathcal{E}}de}{dt} = -\frac{1}{2}U^2\sigma(h) \sin 2\gamma \cos\eta \tag{13}$$

This is the result taken as the starting point of the analysis in the main text, where for simplicity we drop the preceding superscript notation denoting that the derivative is taken in $\mathcal{F}_{\mathcal{E}}$.

**Model of unsteady banked turning** In a steady wind field, the time derivative of the wind velocity $\mathbf{W}$ that a bird experiences as a result of its own motion relative to the Earth is:

$$\frac{{}^{\mathcal{E}}d\mathbf{W}}{dt} = {}^{\mathcal{E}}\nabla\mathbf{W}\mathbf{V} \tag{14}$$

where the preceding superscript notation denotes that the derivative is taken with respect to the Earth-fixed frame of reference $\mathcal{F}_{\mathcal{E}}$. Here, ${}^{\mathcal{E}}\nabla\mathbf{W}$ denotes the gradient of the wind velocity, which is the Jacobian matrix of $\mathbf{W}$. Adding ${}^{\mathcal{E}}d\mathbf{U}/dt$ to both sides and making use of Equation 5 we may rewrite Equation 14 as:

$$\frac{{}^{\mathcal{E}}d\mathbf{V}}{dt} = \frac{{}^{\mathcal{E}}d\mathbf{U}}{dt} + {}^{\mathcal{E}}\nabla\mathbf{W}\mathbf{V} \tag{15}$$

All of the equations to this point have been written in the Earth-fixed axis system $\mathcal{F}_{\mathcal{E}}$, but for the purposes of modelling the bird's turning behaviour, we define a set of body axes $\mathcal{F}_{\mathcal{B}} = (G, \hat{\mathbf{x}}_{\mathcal{B}}, \hat{\mathbf{y}}_{\mathcal{B}}, \hat{\mathbf{z}}_{\mathcal{B}})$. Here, $(\hat{\mathbf{x}}_{\mathcal{B}}, \hat{\mathbf{y}}_{\mathcal{B}}, \hat{\mathbf{z}}_{\mathcal{B}})$ is a set of right-handed orthonormal basis vectors with their origin at the bird's centre of mass $G$. The vector $\hat{\mathbf{x}}_{\mathcal{B}}$ denotes the bird's longitudinal body axis, and is aligned with its aerodynamic velocity vector $\mathbf{U}$ at equilibrium; the vector $\hat{\mathbf{z}}_{\mathcal{B}}$ also lies within the bird's plane of symmetry and points ventrally. The orientation of the body axes with respect to the Earth axes is specified by a set of intrinsic Euler angles $(\psi, \theta, \phi)$ defined in a $z$-$y$-$x$ rotation sequence, where $\psi$ is called the yaw angle, $\theta$ is called the pitch angle, and $\phi$ is called the bank angle.

The time derivative of $\mathbf{U}$ in the Earth-fixed axis system is related to the time derivative of $\mathbf{U}$ in the body axis system by the identity ${}^{\mathcal{E}}d\mathbf{U}/dt = {}^{\mathcal{B}}d\mathbf{U}/dt + {}^{\mathcal{E}}\mathbf{\Omega}^{\mathcal{B}} \times \mathbf{U}$, where the preceding superscript denotes the axis system in which the derivative is taken, and where ${}^{\mathcal{E}}\mathbf{\Omega}^{\mathcal{B}}$ is the angular velocity of the body axes relative to the Earth axes. Using Equation 2 to write ${}^{\mathcal{E}}\nabla\mathbf{W}\mathbf{V} = {}^{\mathcal{E}}\nabla\mathbf{W}\mathbf{U} +^{\mathcal{E}} \nabla\mathbf{W}\mathbf{W}$, we may therefore rewrite the bird's inertial acceleration as:

$$\frac{{}^{\mathcal{E}}d\mathbf{V}}{dt} = \frac{{}^{\mathcal{B}}d\mathbf{U}}{dt} + {}^{\mathcal{E}}\mathbf{\Omega}^{\mathcal{B}} \times \mathbf{U} + {}^{\mathcal{E}}\nabla\mathbf{W}\mathbf{U} +^{\mathcal{E}} \nabla\mathbf{W}\mathbf{W}. \tag{16}$$

We will restrict further analysis to the special case of a steady vertical wind gradient in which the wind is constrained to the horizontal plane and has the same direction everywhere. If we further specify that the otherwise arbitrary horizontal basis vector of the Earth axes, $\hat{\mathbf{x}}_{\mathcal{E}}$, is aligned with the wind, then the Jacobian becomes:

$${}^{\mathcal{E}}\nabla\mathbf{W} = \begin{bmatrix} 0 & 0 & -\sigma(h) \\ 0 & 0 & 0 \\ 0 & 0 & 0 \end{bmatrix} \tag{17}$$

where $\sigma(h) = -\partial W/\partial z_{\mathcal{E}}$ denotes the vertical wind shear gradient with respect to height $h$ above the surface. Given our assumptions on the wind, the ${}^{\mathcal{E}}\nabla\mathbf{W}\mathbf{W}$ term in Equation 16 therefore vanishes, because:

$$
{}^{\mathcal{E}}\nabla\mathbf{W}\mathbf{W} = \begin{bmatrix} 0 & 0 & -\sigma(h) \\ 0 & 0 & 0 \\ 0 & 0 & 0 \end{bmatrix} \begin{bmatrix} W(h) \\ 0 \\ 0 \end{bmatrix} = 0. \tag{18}
$$

where $W(h)$ is the wind speed at height $h$ above the surface. Rewriting Equation 16 therefore yields the equation:

$$
\frac{{}^{\mathcal{E}}d\mathbf{V}}{dt} = \frac{{}^{\mathcal{B}}d\mathbf{U}}{dt} + {}^{\mathcal{E}}\Omega^{\mathcal{B}} \times \mathbf{U} + {}^{\mathcal{E}}\nabla\mathbf{W}\mathbf{U} \tag{19}
$$

describing the bird's acceleration in an inertial frame of reference for a steady horizontal wind field of uniform direction with vertical wind gradient.

Having defined the frame of reference in which each derivative is taken, we are free to resolve the vector components of Equation 19 in whatever axis system we choose. It will prove convenient to do so in the body axes defined by $\mathcal{F}_{\mathcal{B}}$. We write this as:

$$
\left[\frac{{}^{\mathcal{E}}d\mathbf{V}}{dt}\right]_{\mathcal{B}} = \left[\frac{{}^{\mathcal{B}}d\mathbf{U}}{dt}\right]_{\mathcal{B}} + \left[{}^{\mathcal{E}}\Omega^{\mathcal{B}} \times \mathbf{U}\right]_{\mathcal{B}} + \left[{}^{\mathcal{E}}\nabla\mathbf{W}\mathbf{U}\right]_{\mathcal{B}} \tag{20}
$$

where the subscript notation indicates the axes into which the quantities contained in square brackets are projected. We now make the key simplifying assumption that the bird always flies in a symmetric flight condition with its aerodynamic velocity vector $\mathbf{U}$ aligned with its longitudinal body axis $\hat{\mathbf{x}}_{\mathcal{B}}$. This amounts to assuming that the bird is perfectly stable in pitch and yaw, or to assuming that it coordinates its turning to achieve the same effect. With this simplifying assumption, we may write out Equation 20 as:

$$
\left[\frac{{}^{\mathcal{E}}d\mathbf{V}}{dt}\right]_{\mathcal{B}} = \begin{bmatrix} \dot{U} \\ 0 \\ 0 \end{bmatrix} + \begin{bmatrix} 0 & -r & q \\ r & 0 & -p \\ -q & p & 0 \end{bmatrix} \begin{bmatrix} U \\ 0 \\ 0 \end{bmatrix} + \mathbb{T}_{BE} \begin{bmatrix} 0 & 0 & -\sigma(h) \\ 0 & 0 & 0 \\ 0 & 0 & 0 \end{bmatrix} \mathbb{T}_{EB} \begin{bmatrix} U \\ 0 \\ 0 \end{bmatrix} \tag{21}
$$

where $\dot{U}$ is the rate of change of the bird's airspeed $U = \|\mathbf{U}\|$, where $(p, q, r)$ are the bird's angular velocity components about $(\hat{\mathbf{x}}_{\mathcal{B}}, \hat{\mathbf{y}}_{\mathcal{B}}, \hat{\mathbf{z}}_{\mathcal{B}})$, and where $\mathbb{T}_{BE} = [\mathbb{T}_{EB}]^T$ is the change-of-basis matrix transforming from the Earth axes to the body axes:

$$\mathbb{T}_{BE} = \begin{bmatrix} \cos\psi\cos\theta & \sin\psi\cos\theta & -\sin\theta \\ -\sin\psi\cos\phi + \cos\psi\sin\theta\sin\phi & \cos\psi\cos\phi + \sin\psi\sin\theta\sin\phi & \cos\theta\sin\phi \\ \sin\psi\sin\phi + \cos\psi\sin\theta\cos\phi & -\sin\phi\cos\psi + \sin\psi\sin\theta\cos\phi & \cos\theta\cos\phi \end{bmatrix}. \tag{22}$$

Finally, after completing the matrix multiplications and simplifying, Equation 21 can be rewritten:

$$\left[\frac{{}^{\mathcal{E}}d\mathbf{V}}{dt}\right]_{\mathcal{B}} = \begin{bmatrix} \dot{U} \\ 0 \\ 0 \end{bmatrix} + U \begin{bmatrix} 0 \\ r \\ -q \end{bmatrix} - U\sigma(h) \begin{bmatrix} -\cos\psi\sin\theta\cos\theta \\ \sin\psi\sin\theta\cos\phi - \cos\psi\sin^2\theta\sin\phi \\ -\sin\psi\sin\theta\sin\phi - \cos\psi\sin^2\theta\cos\phi \end{bmatrix} \tag{23}$$

which represents the bird's acceleration in an inertial frame of reference, projected onto its body axes. It remains to relate this acceleration to the forces that produce it, through a straightforward application of Newton's second law.

Because the bird's air velocity vector $\mathbf{U}$ is assumed to be aligned with $\hat{\mathbf{x}}_{\mathcal{B}}$, the net thrust-drag force $\mathbf{T} - \mathbf{D}$ acts along $\hat{\mathbf{x}}_{\mathcal{B}}$, whilst the lift force $\mathbf{L}$ acts opposite to $\hat{\mathbf{z}}_{\mathcal{B}}$. The total external force ($\mathbf{F}$) is therefore:

$$[\mathbf{F}]_{\mathcal{B}} = \begin{bmatrix} T - D - mg\sin\theta \\ mg\cos\theta\sin\phi \\ mg\cos\theta\cos\phi - L \end{bmatrix} \tag{24}$$

when resolved in the body axes. Using Newton's second law to relate Equations 23 and 24 as $[\mathbf{F}]_{\mathcal{B}} = m\left[{}^{\mathcal{E}}d\mathbf{V}/dt\right]_{\mathcal{B}}$, we therefore have the component equations:

$$T - D - mg\sin\theta = m\dot{U} + mU\sigma(h)\cos\psi\sin\theta\cos\theta \tag{25}$$

$$mg\cos\theta\sin\phi = mUr - mU\sigma(h)\sin\psi\sin\theta\cos\phi + mU\sigma(h)\cos\psi\sin^2\theta\sin\phi \tag{26}$$

$$mg\cos\theta\cos\phi - L = -mUq + mU\sigma(h)\sin\psi\sin\theta\sin\phi + mU\sigma(h)\cos\psi\sin^2\theta\cos\phi \tag{27}$$

describing the dynamics of flight in a steady shear layer for a bird that is perfectly stable in pitch and yaw. Rearranging equations 26 and 27 we arrive at the following expressions for the angular velocity components:

$$q = \frac{L}{mU} - \frac{g\cos\theta\cos\phi}{U} + \sigma(h)\sin\psi\sin\theta\sin\phi + \sigma(h)\cos\psi\sin^2\theta\cos\phi \tag{28}$$
$$r = \frac{g\cos\theta\sin\phi}{U} + \sigma(h)\sin\psi\sin\theta\cos\phi - \sigma(h)\cos\psi\sin^2\theta\sin\phi \tag{29}$$

The rate of change in the bird's yaw angle $\psi$ is kinematically related to its angular velocity as:

$$\dot{\psi} = \frac{q\sin\phi + r\cos\phi}{\cos\theta}. \tag{30}$$

Substituting Equations 29–28 into Equation 30 and simplifying finally yields the general equation:

$$\dot{\psi} = \frac{L\sin\phi}{mU\cos\theta} + \sigma(h)\sin\psi\tan\theta \tag{31}$$

describing coordinated turning in a shear layer.

The first term on the right hand side of Equation 31 captures how the bird's bank ($\phi$) and pitch ($\theta$) changes the horizontal direction of its longitudinal body axis. The second term captures the aerodynamic effect of the shear layer on the bird's turning, which arises because of our simplifying assumption that the bird is perfectly stable in pitch and yaw, or coordinates its flight to be so. This assumption implies that any sideslip that would otherwise arise as a result of the change in wind speed that the bird experiences as it ascends or descends through the shear layer is immediately trimmed out by weathercocking. It is clear by inspection that the effect of weathercocking is expected to be negligible if the bird ascends or descends through the shear layer in a headwind or tailwind direction, such that $\sin\psi$ is small, but that the effect of weathercocking could be significant otherwise.

The aerodynamic lift $L$ on the wings is given by the lift equation $L = \rho U^2 SC_L/2$, where $\rho$ is air density, $S$ is wing area, and $C_L$ is the lift coefficient, which is a function of the aerodynamic angle of attack. However, as the bird's air velocity vector $\mathbf{U}$ is assumed to coincide with its longitudinal body axis $\hat{\mathbf{x}}_\mathcal{B}$, the product $SC_L$ will remain constant if we assume that the bird holds its wings fixed. Defining a reference airspeed $U_0$ as the airspeed associated with steady level flight at lift equal to body weight, it follows that the lift at airspeed $U$ is $L = mgU^2/U_0^2$. We may therefore rewrite Equation (31) as:

$$\dot{\psi} = \frac{g \sin\phi (1 + \Delta U/U_0)}{U_0 \cos\theta} + \sigma(h) \sin\psi \tan\theta \tag{32}$$

where $U = U_0 + \Delta U$. Assuming that $\Delta U$ is small in comparison with $U_0$ such that $\frac{\Delta U}{U_0} \ll 1$, which is reasonable on the basis that any excess aerodynamic kinetic energy is expected to be converted to gravitational potential energy, it follows that we may approximate the bird's horizontal turn rate as:

$$\dot{\psi} \approx \frac{g \sin\phi}{U_0 \cos\theta} + \sigma(h) \sin\psi \tan\theta \tag{33}$$

which is the same approximation obtained directly by assuming constant airspeed $U = U_0$, and constant lift equal to body weight $L = mg$ in Eq. 31.

The local shear gradient $\sigma(h)$ that a bird experiences is exceedingly difficult to measure empirically. Furthermore, as this term arises through weathercocking, it is heavily dependent on the yaw stability of the bird, which is usually unknown. We have already established that its effect on the bird's turning and will be small if the bird transits the shear layer whilst heading with or against the wind. These are the same flight directions as maximize the instantaneous rate of energy harvesting in dynamic soaring (see Eq. 13). Hence, as a first approximation applicable to birds harvesting energy efficiently from the shear layer, we may adopt the simpler model:

$$\dot{\psi} \approx \frac{g \sin\phi}{U_0 \cos\theta} \tag{34}$$

which is the model of unsteady banked turning used in the main text. More generally, Equation 34 provides a reasonable approximation of turn rate if:

$$\frac{\| \sin\phi \|}{\| \sin\theta \sin\psi \|} \gg \sigma(h) \frac{U_0}{g} \tag{35}$$

where we have made use of the trigonometric identity $\tan\theta = \sin\theta / \cos\theta$.
